# Molecular Disambiguation of Heart Rate Control by the Nucleus Ambiguus

**DOI:** 10.1101/2023.12.16.571991

**Authors:** Maira Jalil, Mary Katherine MacMillan, Veronica A. Gutierrez, Maisie E. Crook, Tatiana C. Coverdell, Ashlyn Dever, Katie Shouse Cox, Daniel S. Stornetta, Yoko B. Wang, Dana C. Schwalbe, Adriana Singh, Lane VanderVoort, Carie R. Boychuk, Stephen B. G. Abbott, John N. Campbell

**Author notes:** **Correspondence to:** John N. Campbell (lead contact), Department of Biology, University of Virginia, 485 McCormick Road, PO BOX 4000328, Charlottesville, VA 22904;, and Stephen B. G. Abbott, Department of Pharmacology, University of Virginia, 1340 Jefferson Park Avenue, Charlottesville, VA 22908. equally contributing senior author (Stephen B. G. Abbott, John N. Campbell).

## Abstract

The nucleus ambiguus (nAmb) provides parasympathetic control of cardiorespiratory functions as well as motor control of the upper airways and esophagus. A subset of nAmb neurons innervates the heart through the vagus nerve to control cardiac function at rest and during key autonomic reflexes such as the mammalian diving reflex. Yet, how these cardiovagal nAmb neurons differ from other nAmb neurons in the adult brain remains unclear. We therefore classified adult mouse nAmb neurons molecularly, anatomically, and functionally. First, our integrated analysis of single-nucleus RNA-sequencing data predicted multiple molecular subtypes of nAmb neurons. Mapping the axon projections of one nAmb neuron subtype, *Npy2r*-expressing nAmb neurons, showed that they innervate cardiac ganglia but not the upper airways or esophagus. Chemogenetically stimulating *Npy2r*+ nAmb neurons robustly decreased heart rate through peripheral muscarinic acetylcholine receptors. Finally, *Npy2r*+ nAmb neurons are activated during voluntary underwater diving, consistent with a cardiovagal function for this nAmb subtype. These results together reveal the molecular organization of nAmb neurons and its control of heart rate.

## INTRODUCTION

The parasympathetic nervous system powerfully slows heart rate by signaling through the vagus nerve. This was famously demonstrated in the mid-19^th^ century by the Weber brothers, who found that electrically stimulating the vagus nerve slowed, and in some cases stopped, the hearts of frogs and mammals ^1,2^. Tonic cardiovagal signaling sets resting HR and mediates autonomic reflexes such as the baroreflex, respiratory sinus arrhythmia, and mammalian diving reflex ^3–6^. The diving reflex, for instance, first described by Edmund Goodwyn in 1786 ^7,8^, is one of the most robust and evolutionarily conserved of autonomic reflexes ^6,9,10^. This reflex occurs when holding breath underwater and triggers a dramatic decrease in heart rate (bradycardia), likely to conserve oxygen stores during an underwater dive ^11^.

Cardiovagal fibers originate primarily from two medullary regions, the nucleus ambiguus (nAmb) and the dorsal motor nucleus of the vagus (DMV) ^12–14^. Cardiovagal axons project through the vagus nerve to synapse on postganglionic parasympathetic neurons in cardiac ganglia ^15,16^. At these ganglionic synapses, cardiovagal release of acetylcholine (ACh) activates cardiac neurons via α4β2 nicotinic ACh receptors ^17^ while potentially suppressing sympathetic input to the same neurons ^18^. CVNs may release other signaling molecules as well – for instance, neuropeptides such as PACAP (gene, *Adcyap1*) to modulate cardiac neuron excitability ^19,20^ and vasoactive intestinal peptide (VIP; gene, *Vip*) to control coronary artery flow ^21,22^. In turn, cardiac neurons synapse on cardiomyocytes in the sinoatrial (SA) and atrioventricular (AV) nodes, where their release of acetylcholine (ACh) on muscarinic acetylcholine receptors slows cardiomyocyte electrical conduction and consequently decreases heart rate, AV conduction, and ventricular contractility. Vagotomy or blocking muscarinic receptors increases heart rate, indicating that parasympathetic tone sets heart rate at rest ^23–25^.

Previous studies suggest the nAmb plays a larger role than the DMV in setting heart rate at rest and during reflexes such as the diving reflex ^26–31^. However, the nAmb is a functionally heterogeneous region which, in addition to its cardiovagal neurons, also contains respiratory parasympathetic neurons and motor neurons for the upper airways and esophagus ^5,32–35^.

Recent findings have raised the possibility that the nAmb’s functional roles are delegated to different molecular subtypes of nAmb neurons ^5,36,37^. Yet, a comprehensive understanding of these molecular subtypes in the adult brain and their anatomical and functional differences is still lacking. We used single-cell transcriptomics and anatomical tracing to define nAmb neuron subtypes and identified one that innervates cardiac ganglia, decreases heart rate upon activation, and is engaged during the diving reflex.

## RESULTS

### Molecular Classification of Nucleus Ambiguus Neurons in the Adult Mouse

To classify molecular subtypes of nucleus ambiguus (nAmb) neurons in adult mice, we compared their genome-wide RNA expression using single-nuclei RNA sequencing (snRNA-seq). First, to enrich for nAmb neurons, we fluorescently labeled their cell nuclei based on expression of the cholinergic marker gene, *Chat,* which encodes choline acetyltransferase (ChAT) and which all nAmb vagal efferents and few neighboring cells express ^36^ (Supplemental Figure 1A). Specifically, we crossed mice in which Cre recombinase expression is driven by the *Chat* gene (Chat-Cre) ^38^, to mice which Cre-dependently express a nuclear-localized mCherry fluorescent protein (H2b-mCherry; “H2b-TRAP” mice). The resulting Chat-Cre::H2b-TRAP mice exhibited H2b-mCherry expression in all peripherally projecting nAmb neurons ^36^ (Supplemental Figure 1B). As a complementary approach, to avoid any cells labeled due to Chat-Cre expression only during development, we injected the nAmb of other adult Chat-Cre mice with an adeno-associated virus (AAV) that Cre-dependently expresses H2b-mCherry. We then isolated mCherry+ cell nuclei from dissected nAmb tissue samples, processed their poly-adenylated RNA into sequencing libraries, and sequenced them (Figure 1A).

**Figure 1:**
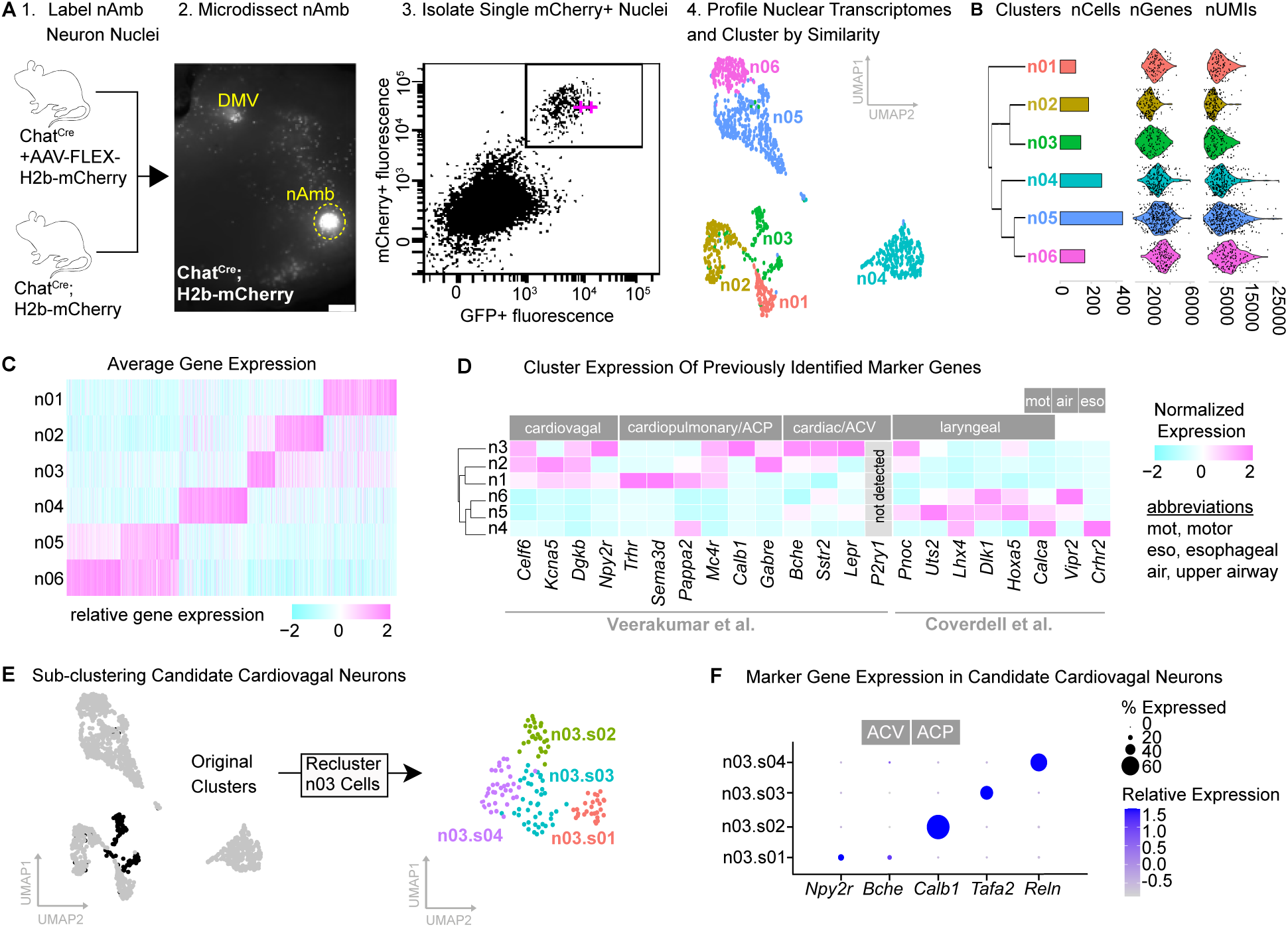
Molecular Classification of Nucleus Ambiguus Neurons in the Adult Mouse. A. Schematic of the single-nuclei RNA-sequencing workflow. From left to right: labeling and isolation of nucleus ambiguus (nAmb) neurons using Chat-Cre;H2b-TRAP mice and Chat-Cre mice injected with a Cre-dependent AAV-H2b-mCherry; selection of mCherry+/GFP+ nuclei from Chat-Cre;H2b-TRAP mice during fluorescence activated nuclei sorting; uniform manifold approximation and projection (UMAP) of ventrolateral medulla vagal neuron clusters. B. From left to right: dendrogram illustrating relatedness of clusters; number of cells, genes and unique molecular identifiers (UMIs; ∼unique RNA transcripts). C. All genes differentially expressed among clusters to a statistically significant extent based on a Bonferroni-adjusted p-value < 0.05. D. nAmb neuron cluster expression of subtype marker genes reported in previously published studies. E. Subclustering cells from candidate cardiovagal cluster n03. F. Expression of candidate marker genes for n03 subclusters. ACP and ACV marker genes labeled based on Veerakumar *et al*.

Our initial dataset contained 3,490 nuclear transcriptomes (“cells”) across three batches (Supplemental Figure 1C). Filtering out low-quality transcriptomes and likely cell doublets left 3,461 cells for further analysis. Grouping the pass-filter cells by their expression of high-variance genes yielded 18 clusters (annotated as c1-c18), each containing cells from each sample batch (Supplemental Figure 1D-G). To identify the nAmb neuron clusters, we plotted expression of positive marker genes associated with cholinergic and motor neurons (vAChT/*Slc18a3*, *Isl1*, *Chat*) and negative marker genes (*Slc32a1*, *Slc17a6*) which are expressed around but not in the nAmb ^36,39–41^ (Supplemental Figure 1H). Based on our results, we annotated five clusters, containing 1,245 neurons in total, as candidate subtypes of nAmb neurons (Supplemental Figure 1H).

We further explored the molecular heterogeneity of nAmb neurons by re-clustering them apart from non-nAmb neurons. This resulted in six transcriptionally distinct clusters (n1-n6) in which we detected on average (+/− standard deviation, S.D.) 2,927 +/− 967 genes per neuron, based on 6,666 +/− 3,565 unique transcripts per neuron (Figure 1B). Each cluster contained cells from each sample batch (Supplemental Figure 1I-J). Comparing the neuron clusters revealed many differentially expressed genes (Figure 1C; Supplemental Figure 1K). Among these was the gene *Adcyap1*, which is enriched in clusters n02 and n03 (Supplemental Figure 1K) and encodes pituitary adenylate cyclase activating polypeptide (PACAP), a neuropeptide found in cholinergic axons innervating the heart in guinea pigs ^19^. These results suggest that clusters n02 and n03 may contain cardiovagal neurons.

We then examined the expression of marker genes reported in previous studies. One of these studies, Coverdell et al., 2022, identified three molecular subtypes among 141 adult mouse nAmb neurons: Crhr2^nAmb^ neurons, which control esophageal motor function; Vipr2^nAmb^ neurons, which innervate the pharynx and larynx; and Adcyap1^nAmb^ neurons, whose anatomy and function were unknown ^36^. Another study, Veerakumar et al. 2022, characterized three molecular subtypes from 203 neonatal mouse nAmb neurons innervating the larynx or heart: ACP neurons and ACV neurons, which can alter cardiopulmonary and cardiovascular function, respectively, and larynx-projecting neurons ^37^. When we plotted the expression of these published marker genes in our dataset, we found that clusters n01, n02 and n03 likely correspond to cardiovagal parasympathetic motor neurons, whereas clusters n04, n05 and n06 aligned to branchial motor neurons that innervate skeletal muscles of the upper airway and esophagus (Figure 1D).

Cluster n03 expressed markers of two previously defined subtypes of cardiovagal neurons, ACP neurons and ACV neurons (Figure 1D), suggesting heterogeneity within this cluster. Re-clustering only the neurons from n03 resolved four sub-clusters (n03.s01-04; Figure 1E). One of these sub-clsuters expressed a marker of ACV neurons (*Bche*), while another expressed a marker of ACP neurons (*Calb1*; Figure 1F). Interestingly, the putative ACV neurons also expressed *Npy2r*, the gene encoding the neuropeptide Y2 receptor, which modulates vagal control of heart rate ^42^ and is also expressed by vagal sensory neurons that control heart rate ^42–44^. Together, our results identify five candidate molecular subtypes of neurons in the adult nAmb and raise the possibility that *Npy2r⁺* nAmb neurons contribute to heart-rate control.

To compare our nAmb neuron subtypes to those identified by previous studies, we mapped single-cell transcriptomes from the Coverdell et al. and Veerakumar et al. studies onto our nAmb neuron clusters using reference query mapping. This method projects query cells onto reference cell clusters without altering the reference data structure ^45^. Interestingly, while the Coverdell et al. cell clusters largely mapped to different reference cell clusters (Supplemental Figure 2A) as expected, the Veerakumar et al. cells subdivided among the same three reference clusters (Supplemental Figure 2B). This was surprising, since, for instance, the larynx-projecting subtypes in Coverdell et al. (Vipr2^nAmb^ neurons) and Veerakumar *et al.* (laryngeal neurons) mapped to different reference cell clusters. When we assessed the confidence of cell mapping, most Coverdell *et al.* cells mapped with high confidence (>0.8; Supplemental Figure 2C) whereas most Veerakumar *et al.* cells mapped with lower confidence (<0.6; Supplemental Figure 2D). This suggests that the adult nAmb neurons in Coverdell *et al*. are more like the adult nAmb neurons in our present dataset than are the neonatal nAmb neurons in Veerakumar *et al*., potentially due to developmental differences in gene expression. In line with this possibility, the larynx-projecting neurons of Veerakumar *et al*. but not in those of Coverdell et al. (Supplemental Figure 2E-F) expressed the gene *Adcyap1*, consistent with the broader expression of *Adcyap1* in the developing brain ^46^. Thus, the ability to match neonatal and adult nAmb neurons with the reference query approach may be limited by developmental differences in gene expression.

### Partially Overlapping Populations of Nucleus Ambiguus Neurons Express Npy2r and Adcyap1

Given the likelihood that cardiovagal nAmb neurons express *Npy2r* and *Adcyap1*, we mapped the transcripts of these genes in the adult mouse nAmb using RNA fluorescence *in situ* hybridization (RNA FISH). First, to label peripherally-projecting neurons, including all nAmb vagal efferent neurons, we intraperitoneally injected adult C57BL6/j mice with the retrograde tracer Fluorogold ^47,48^. We then performed RNA FISH for *Adcyap1*, *Npy2r*, and *Chat* transcripts in coronal sections and imaged them throughout the nAmb’s compact, semi-compact, and loose subregions. We observed *Npy2r*+ nAmb vagal efferent neurons scattered throughout these subregions (representative images in Figure 2A), consistent with the known anatomical distribution of cardiovagal nAmb neurons ^13,30,37,49^. Quantifying nAmb neurons expressing *Adcyap1* and/or *Npy2r* revealed a partially overlapping distribution. Specifically, 4% of nAmb neurons expressed both *Npy2r* and *Adcyap1*, 3% expressed *Npy2r* but not *Adcyap1*, and <1% expressed *Adcyap1* but not *Npy2r* (Figure 2A, B). Thus, only a small minority of nAmb vagal efferent neurons express *Adcyap1* and/or *Npy2r*. Of note, the 8% of nAmb neurons expressing *Adcyap1* and/or *Npy2r*, extrapolated for the total number of adult mouse nAmb neurons ^50^, would be approximately 47 neurons per nAmb, slightly below previous estimates of retrogradely-labeled cardiovagal nAmb neurons in mice (50-60 neurons) ^37^.

**Figure 2:**
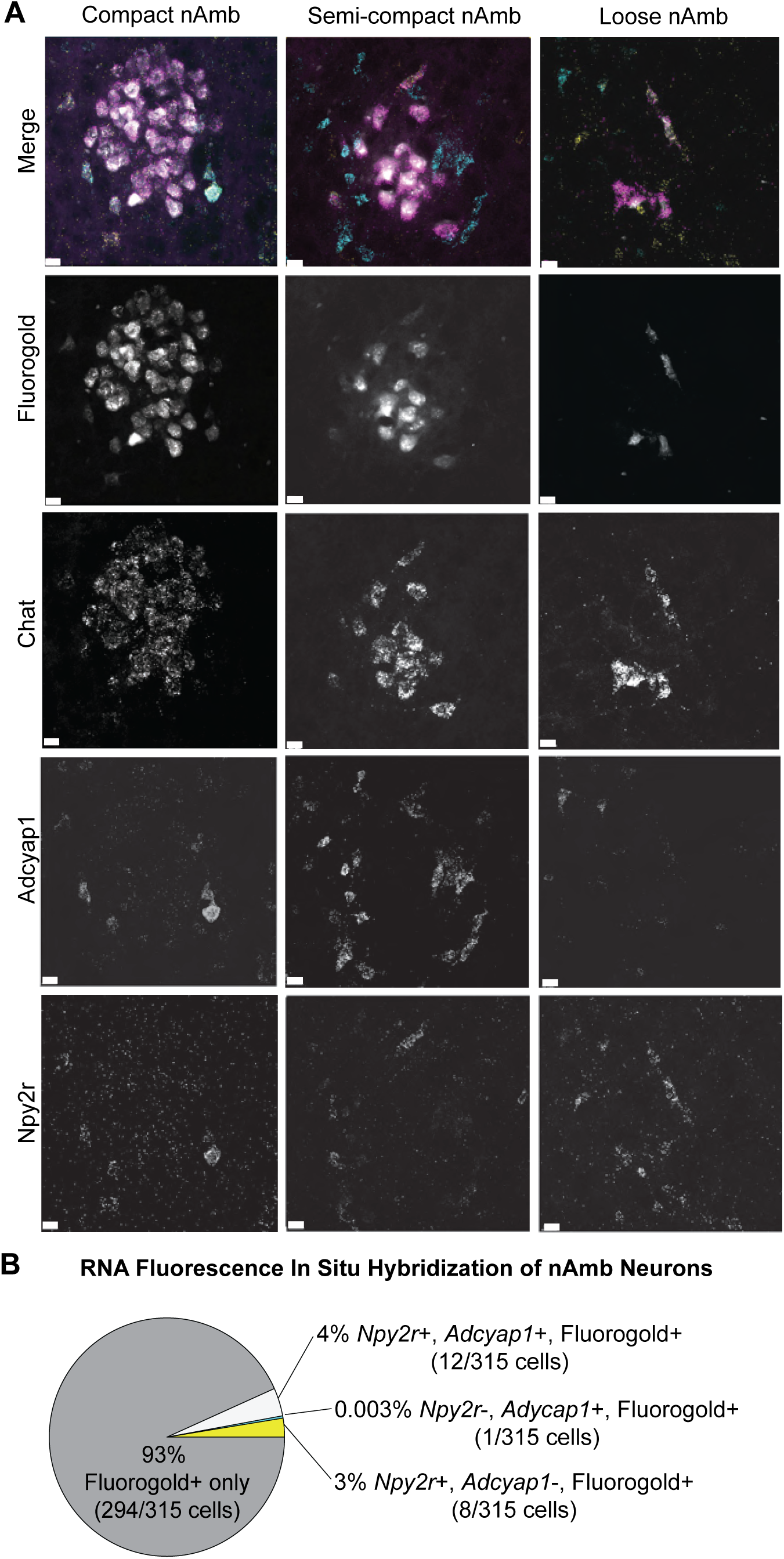
Partially Overlapping Populations of Nucleus Ambiguus Neurons Express *Npy2r* and *Adcyap1*. A. RNAscope fluorescent in situ hybridization (RNA FISH) for *Npy2r* and *Adcyap1* in compact, semi-compact, and loose subdivisions of the nAmb. All nAmb vagal efferent neurons were identified by RNA FISH for *Chat* and systemic Fluorogold labeling (scale bar, 20 μm). All individual color channels are displayed in grayscale. In the merged image, Fluorogold is shown in grey, *Chat* in magenta, *Adcyap1* in cyan, and *Npy2r* in yellow. B. Quantification of RNA FISH data (n=3 mice)

### Retrograde Labeling from the Heart Reveals Cardiovagal Populations in the nAmb and Intermediate Zone

To identify heart-projecting vagal motor neurons, we injected the retrograde tracer Fluorogold into the pericardiac fat pad near the posterior right atrioventricular junction, where cardiovagal nerve endings terminate ^51^ (Figure 3A). One week later, we collected brainstem tissue for Fluorogold immunofluorescence and RNA FISH. On average, we identified 40 ± 24 Fluorogold-labeled neurons per mouse bilaterally in the nAmb, distributed across the full rostral–caudal extent of the nAmb (Figure 3B; n = 3 mice). Among these, 27% contained *Npy2r* transcripts by RNA FISH (Figure 3C). These results indicate that *Npy2r+* nAmb (Npy2r^nAmb^) neurons are a subset of cardiovagal nAmb neurons.

**Figure 3:**
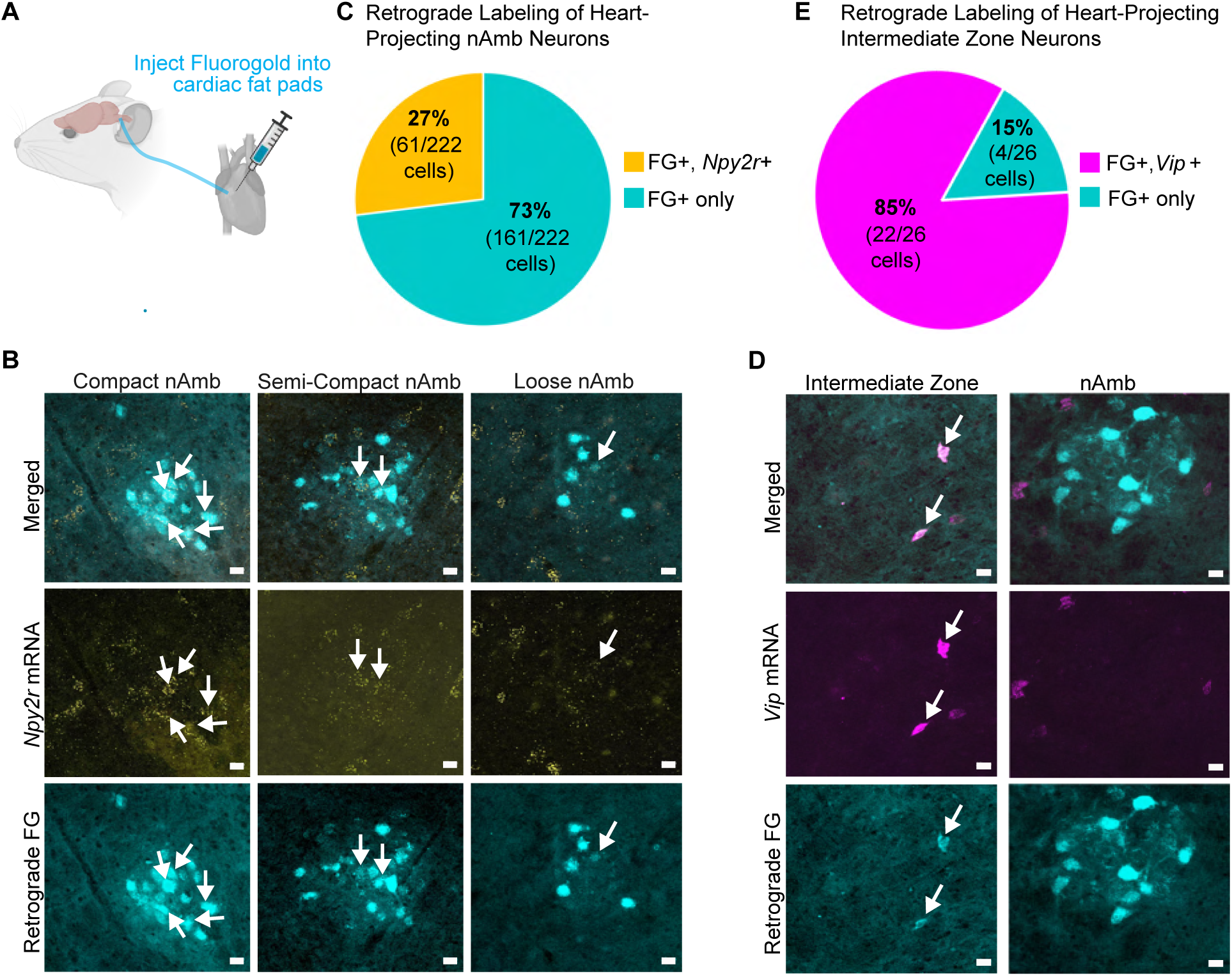
Retrograde Labeling from the Heart Reveals Cardiovagal Populations in the nAmb and Intermediate Zone. A. Schematic of retrograde tracing from heart with Fluorogold. B. *Npy2r* RNA FISH and Fluorogold labeling in the compact, semi-compact and loose nAmb. Scale bar, 20 μm (n=3 mice). C. Quantification of *Npy2r*+ cells as a subset of Fluorogold+ nAmb cells. D. *Vip* RNA FISH and Fluorogold labeling in the compact, semi-compact and loose nAmb. Scale bar, 20 μm (n=3 mice). E. Quantification of *Vip*+ cells labeled with retrograde Fluorogold in the intermediate zone.

Another neuronal cluster identified in our single-nuclei RNA-seq analysis, cluster n02 (*Vip*+/*Chat*+/*Phox2b*+), was molecularly most like Npy2r⁺ nAmb neurons (cluster n03; see cladogram in Figure 1B), raising the possibility that n02 neurons are also cardiovagal. To test this, we retrogradely labeled heart-projecting neurons with intracardiac Fluorogold, as described above, and assessed co-localization of Fluorogold with RNA FISH for Vip, a marker for cluster n02 neurons, in the ventrolateral medulla. Surprisingly, *Vip* expression was absent from Fluorogold-labeled nAmb neurons (Figure 3D). Instead, we observed robust *Vip* expression in Fluorogold-labeled neurons of the intermediate zone between the nAmb and the DMV, where 85% of Fluorogold-labeled neurons also expressed *Vip* (Figure 3D-E). These findings indicate that the *Vip+* neuron cluster (n02) corresponds to cardiovagal neurons of the intermediate zone rather than the nAmb, and that their inclusion in our single-nuclei RNA-seq analysis likely reflects a dissection artifact. Of note, the *Vip* gene product, vasoactive intestinal peptide (VIP), contributes to parasympathetic cardiac regulation by modulating heart rate, influencing intracardiac ganglion excitability, and interacting with cholinergic signaling mechanisms that generate cardioinhibitory output ^52,53^.

Together, our retrograde-tracing results demonstrate that the heart receives vagal motor input from at least two molecularly and anatomically distinct neuron populations: *Npy2r+* neurons in the nAmb and *Vip*+ neurons in the intermediate zone.

### Npy2r^nAmb^ Neurons Innervate Cardiac Ganglia

Since retrograde tracers injected into the heart can spread to other visceral tissues and label non-cardiovagal neurons ^54^, we performed cell-type specific anterograde tracing to confirm that Npy2r^nAmb^ neurons innervate the heart. Specifically, we labeled the axons of nAmb neurons with an adeno-associated virus (AAV) that Cre-dependently expresses the red fluorescent protein, tdTomato (AAV9-DIO-tdTomato). We confirmed the Cre dependency of AAV9-DIO-tdTomato by injecting it into the ventrolateral medulla of adult C57BL6/j mice, where we failed to observe tdTomato immunofluorescence three weeks later at the injection sites (n=3 mice; Supplemental Figure 3A). Next, to target this AAV to Npy2r^nAmb^ neurons, we obtained Npy2r-Cre mice ^44^. We validated the specificity and sensitivity of Cre expression in these mice by co-localizing *Npy2r* and *Cre* transcripts by RNA FISH in the nAmb. In Npy2r-Cre mice, 92% (145 of 157) *Npy2r* mRNA+ neurons were also *Cre* mRNA+, and 100% (145 of 145) of *Cre* mRNA+ cells were also *Npy2r* mRNA+ (n=3 mice; Supplemental Figure 3B). These results validate the specificity of AAV9-DIO-tdTomato and Npy2r-Cre mice for genetically targeting Npy2r^nAmb^ neurons.

To fluorescently label Npy2r^nAmb^ axons for anterograde tracing, we injected the ventrolateral medulla of Npy2r-Cre mice with AAV9-DIO-tdTomato (Figure 4A). As a positive control, we also injected the ventrolateral medulla of Chat-Cre mice ^38^, which express Cre activity in all nAmb vagal efferent neurons ^36^ (Chat^nAmb^ neurons). After waiting at least three weeks for AAV transgene expression, we observed tdTomato immunofluorescence in a subset of nAmb neurons, identified based on their immunofluorescence of the cholinergic marker ChAT (choline acetyltransferase). While some neurons outside the nAmb in Npy2r-Cre mice were also tdTomato+, none were ChAT+ (Figure 4B), indicating that they were not vagal efferents and so would not confound our anterograde tracing results. Importantly, we observed no tdTomato+ neurons in the dorsal motor nucleus of the vagus (DMV), another source of heart innervation (Supplemental Figure 3C) ^13,55^.

**Figure 4:**
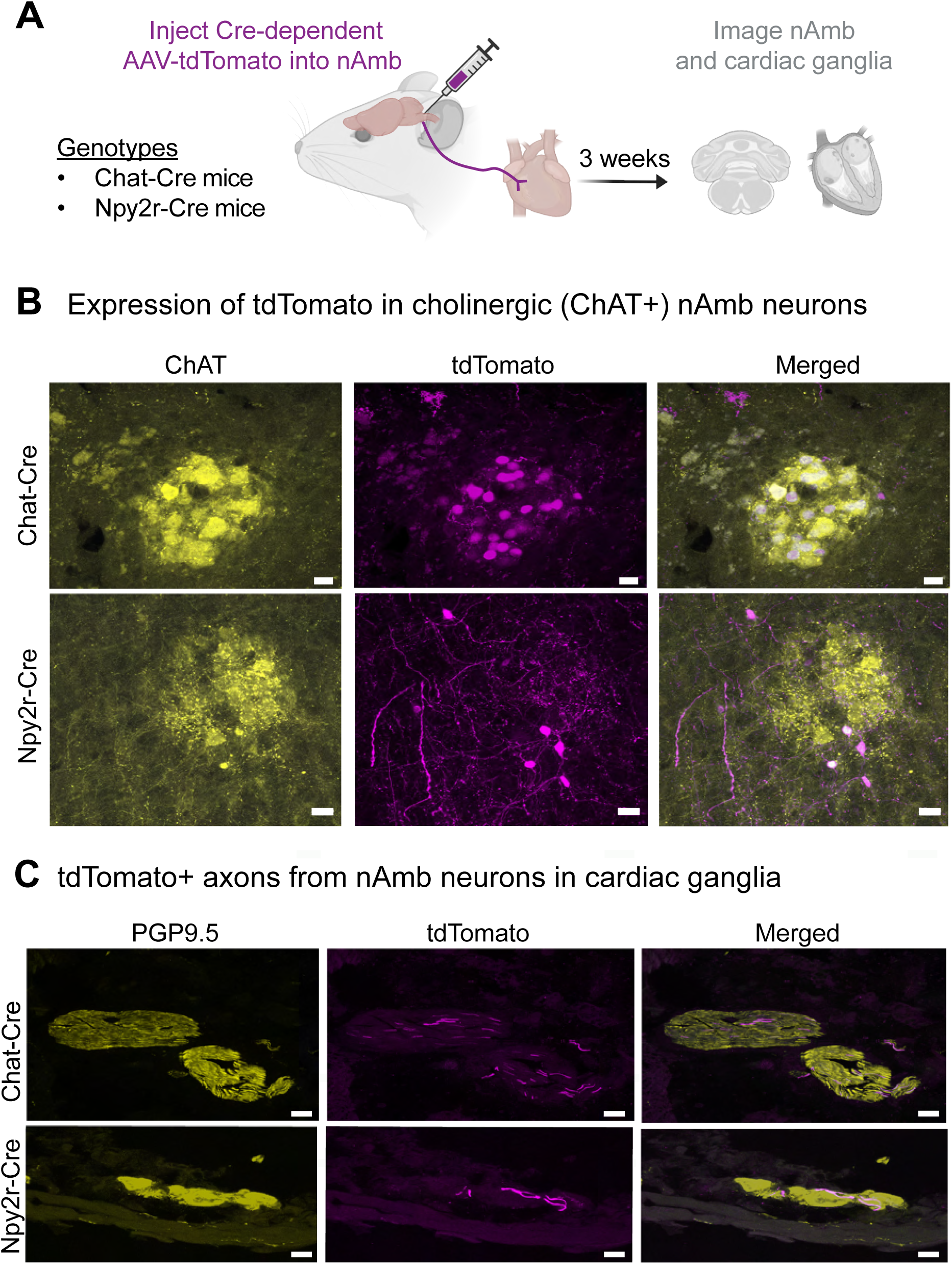
Npy2r^nAmb^ Neurons Innervate Cardiac Ganglia. A. AAV9-FLEX-tdTomato injected into right side nAmb of Npy2r-Cre mice and, as positive controls, Chat-Cre mice (n=3 mice per genotype). Figure panel created with Biorender.com. B. tdTomato and ChAT immunofluorescence in nAmb of Chat-Cre mice and Npy2r-Cre mice. Scale bar, 20 um. C. tdTomato axons of Chat-Cre and Npy2r-Cre nAmb neurons in PGP9.5 immunofluorescent cardiac ganglia. Scale bar, 20 um.

To determine whether Npy2r^nAmb^ neurons innervate the heart, we then imaged immunofluorescence of tdTomato and the neuronal marker protein PGP9.5 in cardiac tissue from the same mice. PGP9.5 is expressed generally by neurons, including all cardiac ganglia neurons in rodents ^56^. We observed fibers immunofluorescent for both tdTomato and PGP9.5 innervating PGP9.5+ cardiac ganglia in all Npy2r-Cre mice as well as positive-control Chat-Cre mice (Figure 4C; n=3 mice per genotype; at least two innervated ganglia per mouse). These results indicate that Npy2r^nAmb^ neurons innervate cardiac ganglia, confirming their identity as cardiovagal neurons.

### Npy2r^nAmb^ Neurons Do Not Innervate the Upper Airways or Esophagus

Since the nAmb also contains motor neurons for the esophagus and upper airways ^33–35,57,58^, we investigated whether Npy2r^nAmb^ neurons project to those tissues using a chromogenic anterograde tracing approach. First, we injected an AAV encoding Cre-dependent placental alkaline phosphatase (AAV9-FLEX-PLAP) into the ventrolateral medulla of Npy2r-Cre mice (n=3 mice). As a positive control, we performed the same injections in Chat-Cre mice (n=3 mice; Figure 5A) ^36,59^. At least 3 weeks later, we collected the medulla and en bloc tissue samples of the esophagus and upper airways for histology. We observed PLAP immunofluorescence in ChAT-positive neurons along the rostrocaudal extent of the nAmb (Figure 5B), but none in the DMV (Supplemental Figure 4), indicating that our injections were restricted to cholinergic nAmb neurons.

**Figure 5:**
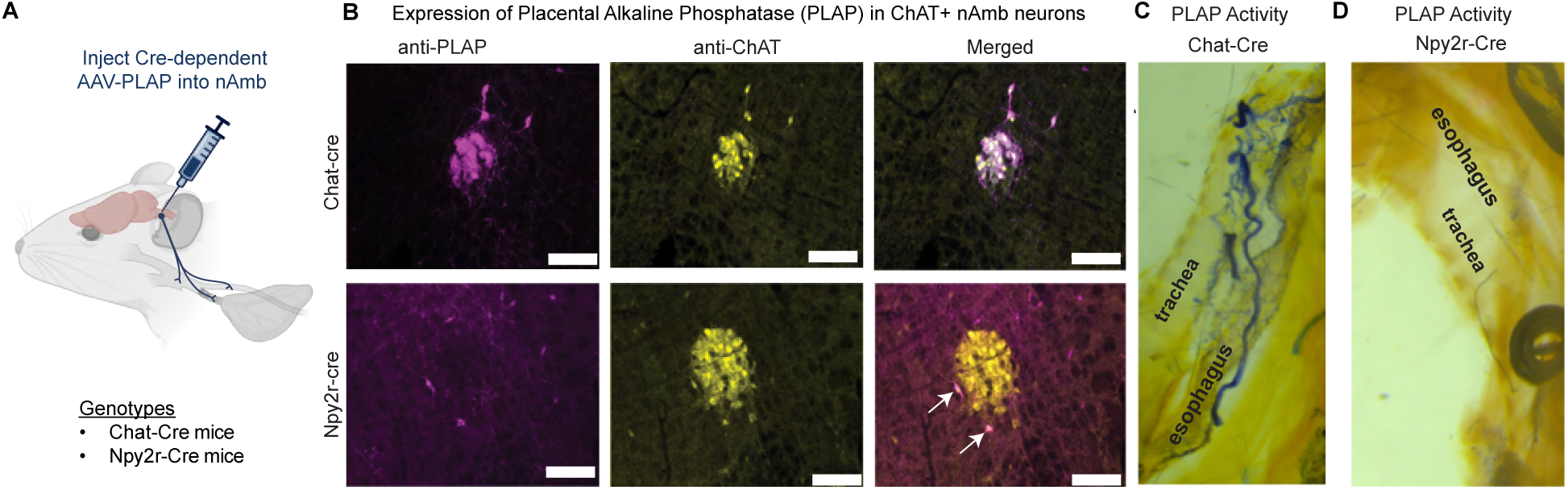
Npy2r^nAmb^ Neurons Do Not Innervate the Upper Airways or Esophagus. A. Strategy for targeting AAV-PLAP expression to Cre-expressing nAmb neurons of Chat-Cre and Npy2r-Cre mice. Figure panel created with Biorender.com. B. PLAP and ChAT immunofluorescence in nAmb cells (n=3 mice). Arrows indicate double-positive cells in an Npy2r-Cre mouse nAmb. Scale bar, 70 um. C. PLAP-stained axons from Chat^nAmb^ neurons in the upper airways and esophagus (n=3 mice). D. PLAP-stained axons from Npy2r^nAmb^ neurons in the upper airways and esophagus (n=3 mice).

To visualize the axons of PLAP+ neurons, we chromogenically stained them while optically clearing the esophagus and upper airways as previously described ^36,60^. Consistent with previous studies, we observed PLAP-stained axons from Chat^nAmb^ neurons throughout the larynx, pharynx, and esophagus (n=3 mice; representative image in Figure 5C) ^36^. However, we did not observe any such projections from Npy2r^nAmb^ neurons (n=3 mice; representative image in Figure 5D). Our anterograde tracing studies together indicate that Npy2r^nAmb^ neurons innervate the heart but not the upper airways or esophagus.

### Chemogenetically Activating Npy2r^nAmb^ Neurons Lowers Heart Rate

Since Npy2r^nAmb^ neurons innervate the heart, we investigated whether they could control heart rate. To selectively target these neurons, we relied on an intersectional chemogenetics approach, using an AAV which Cre- and Flp-dependently expresses the excitatory chemogenetic receptor hM3Dq ^51^. We first validated the specificity of this AAV by injecting it into the ventrolateral medulla of Chat-Cre;Phox2b-Flp mice, targeting nAmb neurons in general by their co-expression of *Chat* and *Phox2b* (Supplemental Figure 5A) ^36^. We then quantified co-localization of immunofluorescence for ChAT and hM3Dq-HA in the nAmb, which is the only region where we observed hM3Dq-HA expression. Our results demonstrate that 91% of hM3Dq-HA+ cells were also ChAT+ in the nAmb, while 9% of hM3Dq-HA+ nAmb cells were ChAT-(n=132 cells in total from 5 mice; Supplemental Figure 5B-C). The small percentage of hM3Dq-HA+/ChAT- cells most likely represents neurons with low ChAT expression, rather than Cre-independent AAV transgene expression, since we failed to detect any hM3Dq-HA immunofluorescence after injecting this AAV into the nAmb of Phox2b-Flp mice that lacked Chat-Cre (Supplemental Figure 5D). These results validate our previous finding that this AAV only expresses hM3Dq-HA in cells that bear both Cre and Flp recombinases ^51^.

Next, to selectively activate Npy2r+ nAmb neurons while measuring heart rate, we injected the Cre- and Flp-dependent AAV-hM3Dq-HA into the ventrolateral medulla of Npy2r-Cre;Chat-Flp;R26^ds-HTB^ mice (Figure 6A). This resulted in expression of hM3Dq-HA expression in a subset of nAmb neurons, mostly located in the external formation, where cardiovagal neurons largely reside (Figure 6B) ^61^. Of note, crossing Npy2r-Cre;Chat-Flp mice to the Cre/Flp reporter mouse strain, R26^ds-HTB^ (H2b-GFP) ^62^ resulted in H2b-GFP expression in the compact nAmb (Figure 6B), where esophageal motor neurons reside ^34–36,63^, which may be due to transient developmental expression of Npy2r-Cre in the compact nAmb (lineage labeling). In contrast, viral labeling of Npy2r-Cre;Chat-Flp neurons with hM3Dq-HA in adult mice mostly labeled neurons in the external nAmb, with relatively few in the compact nAmb, consistent with both their cardiovagal identity and our results described above from virally infecting Npy2r-Cre nAmb neurons in the anterograde tracing studies.

**Figure 6.**
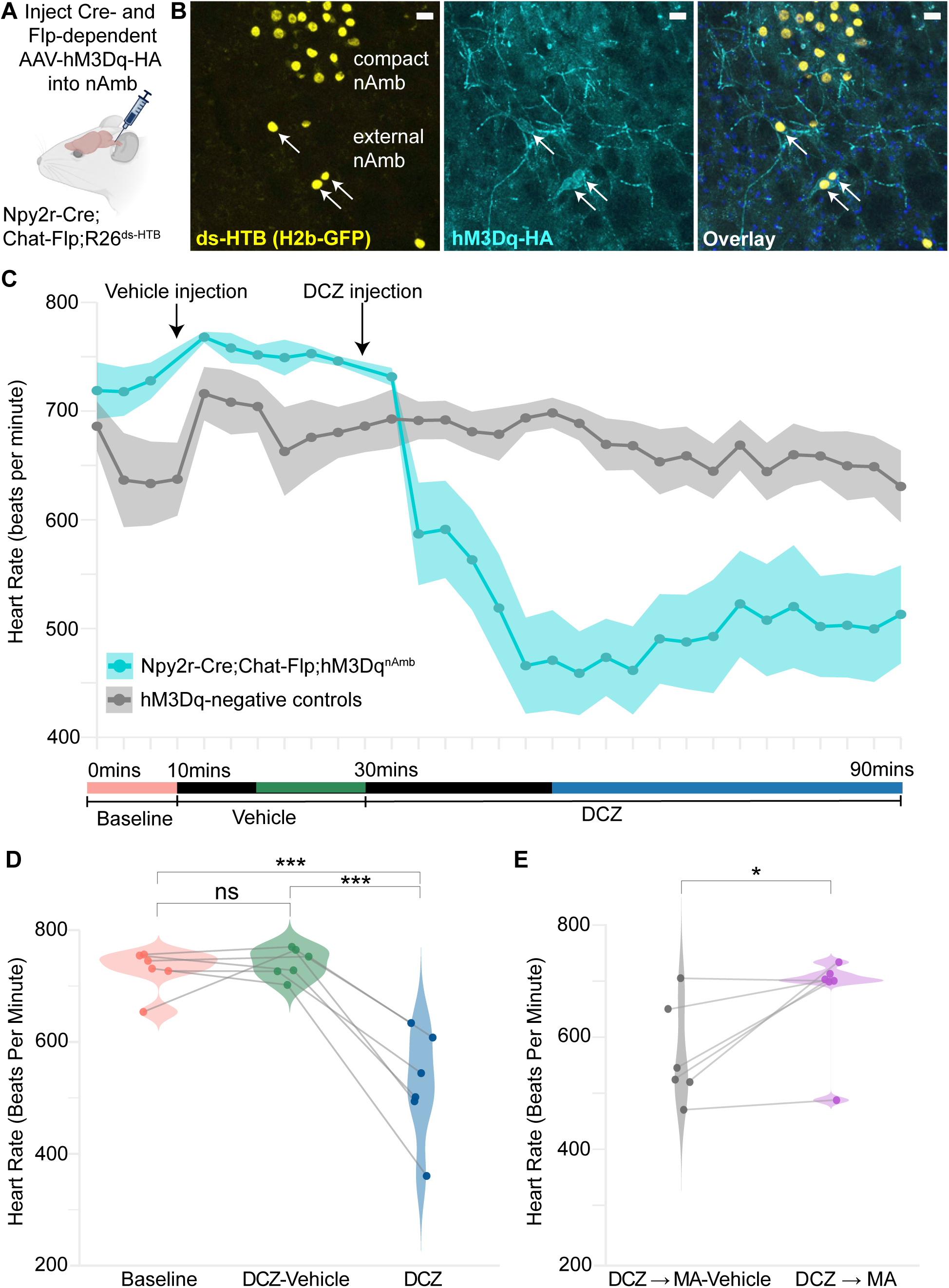
Chemogenetically Activating Npy2r^nAmb^ Neurons Lowers Heart Rate. A. Schematic of experimental design. Figure panel created with Biorender.com. B. Representative image of H2b-GFP (R26^ds-HTB^ reporter) and hM3Dq-HA immunofluorescence in nAmb of Npy2r-Cre;Chat-Flp;R26^ds-HTB^ mice after injecting Cre-and Flp-dependent AAV-hM3Dq-HA into their ventrolateral medulla. Arrows indicate hM3Dq-HA-positive cells. The H2b-GFP observed in the compact nAmb is likely the result of transient developmental expression of Npy2r-Cre. C. Heart rate measured by non-invasive EKG while chemogenetically activating Npy2r^nAmb^ neurons (n=6 mice). Heart rate was recorded every 3 min, with each line representing an individual measurement across time. For statistical analyses, heart-rate values were averaged within each defined time window (baseline, vehicle, DCZ), and these window-averaged values were then used for all comparisons. The color-shaded parts of the timeline below the figure indicate the time windows used for averaging HR. DCZ treatment significantly reduced heart rate relative to both baseline (HR_Baseline_ = 728 ± 16 bpm; mean ± SD) and DCZ-vehicle treatment (HR_vehicle_ = 741 ± 11 bpm; mean ± SD). The reduction was observed for DCZ (HR_DCZ_ = 524 ± 40 bpm; mean ± SD) with an estimated mean difference of 204.35 ± 34.9 bpm compared to baseline (one-way repeated-measures ANOVA with Tukey’s post hoc test, p = 5.8 × 10⁻⁴, t(10) = 5.63) and 217.04 ± 36.3 bpm compared to vehicle (one-way repeated-measures ANOVA with Tukey’s post hoc test, p = 3.6 × 10⁻⁴, t(10) = 5.98). ***, p < 0.001. Colors correspond to the colored parts of the timeline below panel C. D. Administering methyl-atropine bromide (MA; 1.0 mg/kg, i.p.) rescued heart rate (HR_MA_ = 673 ± 37 bpm; mean ± SD), whereas MA-vehicle injections (HR_MA-veh_ = 569 ± 36 bpm; mean ± SD) failed to reverse the bradycardia (estimated mean difference 103.64 ± 38.18 bpm, paired t-test, t(5) = −2.71, df= 5 p = 0.042). *, p < 0.05.

We next measured heart rate (HR) non-invasively in awake mice ^51^ while chemogenetically activating their Npy2r^nAmb^ neurons with a non-metabolizable ligand for hM3Dq, deschloroclozapine (DCZ) ^64^. After acclimation to the ECG system, each mouse underwent a series of intraperitoneal injections while we recorded its HR (n=9 mice). Injecting the vehicle for DCZ into Npy2r-Cre;Chat-Flp;hM3Dq^nAmb^ mice did not significantly affect HR, relative to baseline (HR_Baseline_, 728 ± 16 bpm; HR_veh,_ 741 ± 11 bpm _;_ mean ± S.D.; Tukey post-hoc following one way repeated-measures ANOVA, p= 0.94). Subsequently injecting DCZ (5 mg/kg; i.p.), however, significantly lowered their HR to 524 ± 40 bpm (HR_veh_ vs. HR_DCZ_, one-way repeated measures ANOVA, p = 3.6 × 10⁻⁴; Figure 6C-D). DCZ also significantly lowered HR from baseline levels (HR_baseline_ vs. HR_DCZ_, one-way repeated measures ANOVA, p = 5.8 × 10^−4^; Figure 6D). However, injecting DCZ into wildtype mice did not significantly alter their heart rate relative to either baseline or DCZ vehicle injection (2% DMSO), indicating that DCZ on its own does not affect heart rate (n=6 mice; Figure 6C, Supplemental Figure 5E). Our results thus indicate that activating Npy2r^nAmb^ neurons can robustly decrease heart rate.

The vagus nerve decreases HR through a canonical pathway in which postganglionic cardiac neurons inhibit sinoatrial node myocytes by activating muscarinic acetylcholine receptors. To determine whether Npy2r^nAmb^ neurons use this pathway to decrease HR, we administered methyl-atropine (“MA”; 1.0 mg/kg; i.p.), a peripherally restricted muscarinic receptor antagonist. Mice were administered DCZ and then either MA or its vehicle in separate trials while we recorded their heart rate (n=9 mice). Administering MA after DCZ significantly increased HR (Figure 6E). In addition, HR after MA injection was significantly higher than after MA-vehicle injection (Figure 6E). Thus, activating Npy2r^nAmb^ neurons decreases HR through a canonical cardiovagal signaling pathway involving peripheral muscarinic receptors.

### Voluntary Underwater Diving Activates Npy2r^nAmb^ Neurons

Voluntary underwater submersion elicits the diving reflex in mammals, including a powerful vagal bradycardia ^10^. Based on the bradycardic response we observed when chemogenetically activating Npy2r^nAmb^ neurons, we hypothesized that these neurons would be activated by the diving response. We tested this hypothesis by evaluating expression of the immediate early gene and neuronal activation marker, *Fos*, in nAmb neurons following voluntary diving. To achieve this, we trained mice to voluntarily dive up to 20 cm underwater to reach an escape platform (Figure 7A). We first validated that this model evokes a diving response by measuring ECG with radio-telemetry implants (Figure 7B). Heart rate decreased markedly upon initiation of the dive and remained low while underwater before recovering to baseline, following a brief overshoot after the mice resurfaced (n=10 mice; Figure 7A,B), consistent with previous descriptions. We then performed a session of repeated diving for up to 30 mins to robustly activate the diving response before perfusing mice for histology. Controls were handled and allowed to explore the test chamber with a shallow water level but did not perform any dives (Figure 7C). In mice that performed voluntary diving, we observed a 174% increase in the proportion of Npy2r^nAmb^ neurons that expressed *Fos* (n=6-7 mice; dive vs. control, 44.5 ± 5.5% vs. 16.3 ± 4.3% Npy2r^nAmb^ neurons expressing Fos, unpaired t-test, t=3.915, df=11, p=0.0024) (Figure 7D,F,E). Cell counts for total Npy2r^nAmb^ neurons were similar between cases (42 ± 6 vs. 45 ± 6 cells per unilateral nAmb in a 1 in 3 series, unpaired t-test, t=0.3679, df=11, p=0.72), and a similar proportion of FG cells expressed Npy2r between cases (17.6 ± 1.6 vs. 20.6 ± 3.5% of all FG cells expressed *Npy2r*, unpaired t-test, t=0.7933, df=11, p=0.44). These data indicate that Npy2r^nAmb^ neurons are activated during voluntary diving, consistent with these cells being functionally important for the parasympathetic regulation of the heart.

**Figure 7:**
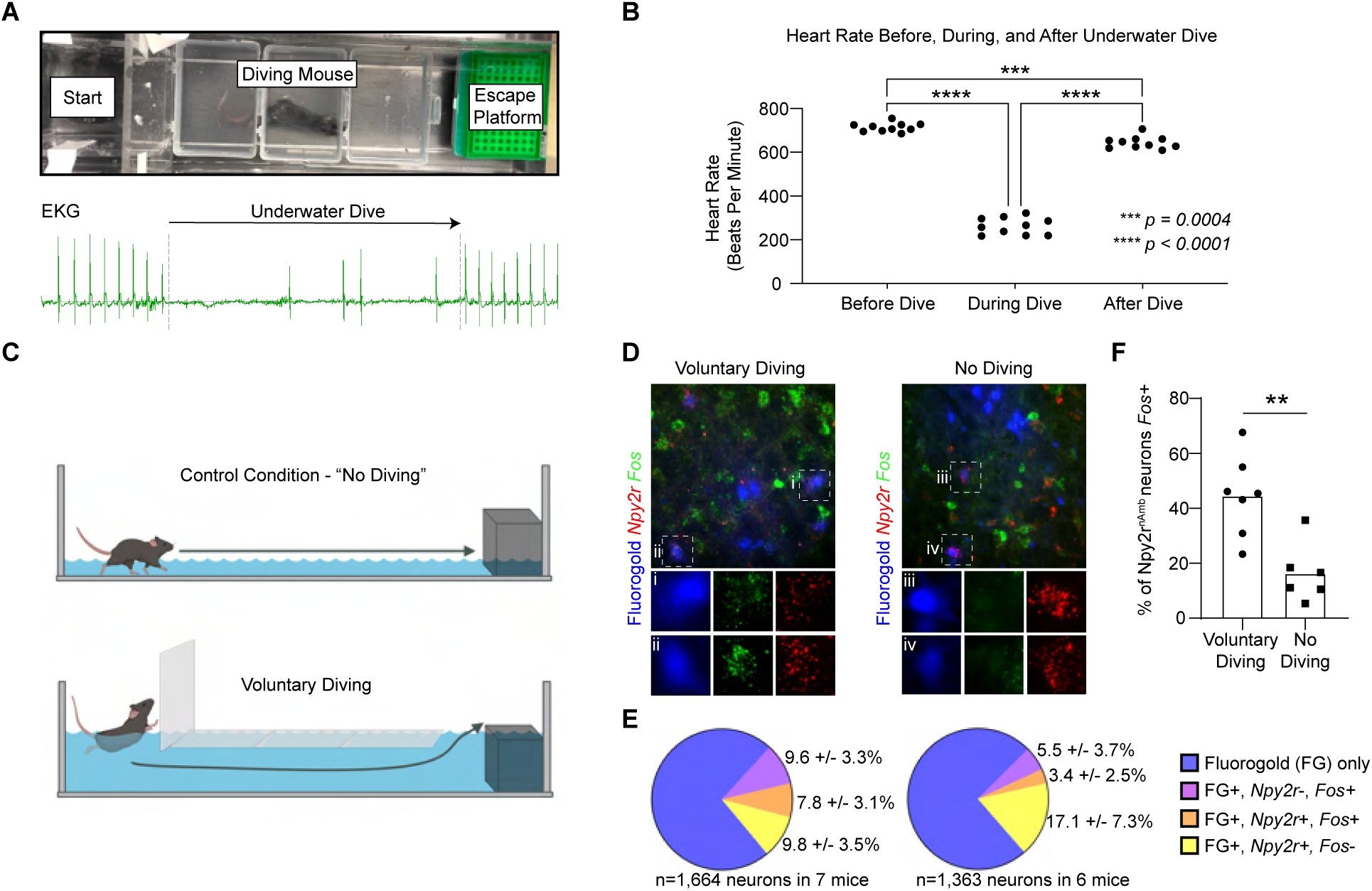
Voluntary Underwater Diving Activates Npy2r-Expressing Nucleus Ambiguus Neurons. A. Experimental set-up for voluntary diving assay (upper panel). EKG recorded during a voluntary dive (lower panel). EKG signal was collected using an implanted radio telemetry device. B. Grouped data for heart rate during voluntary diving. C. Experimental design to compare *Fos* expression in Npy2r nAmb neurons during voluntary diving and no-diving controls. Trained mice performed approximately 20-40 dives over a 20-30 min period followed by 30 min recovery period before perfusion for histology. Mice in the control condition performed no dives but were subjected to similar handling procedures. Figure panel created with Biorender.com. D. RNA FISH of the immediate early gene *Fos* and the nAmb neuron subtype marker *Npy2r* after voluntary diving and control conditions. Systemically administered Fluorogold labels all nAmb vagal efferent neurons. E. Quantification of *Fos* and *Npy2r* expression in Fluorogold+ nAmb neurons during voluntary diving (left) and control (right) conditions. F. Comparison of the proportion of *Npy2r*+ nAmb neurons that expressed *Fos* in the voluntary diving and control condition. Unpaired t-test, t=3.915, df=11, p=0.002

## DISCUSSION

This study identified cardiovagal nAmb neurons based on their molecular, anatomical, and functional features. Transcriptomic profiling and clustering analysis of 1,245 *Chat* expressing neurons in the ventrolateral medulla indicated nine molecular subtypes. RNA fluorescence *in situ* hybridization (RNA FISH) showed that *Adcyap1*+ nAmb neurons and *Npy2r*+ nAmb (Npy2r^nAmb^) neurons are rare and partially overlapping populations of nAmb neurons.

Retrograde and anterograde tracing studies revealed that Npy2r^nAmb^ neurons innervate cardiac ganglia but not the esophagus and airways. Activating Npy2r^nAmb^ neurons with intersectional chemogenetics significantly decreased heart rate through a canonical cardiovagal signaling pathway. Finally, voluntary underwater diving significantly increased the percentage of *Fos*-expressing Npy2r^nAmb^ neurons, indicating that these neurons are activated during the diving reflex. Overall, our study identifies Npy2r^nAmb^ neurons as molecular subtype which selectively innervates cardiac ganglia, is activated during the diving reflex, and decreases heart rate upon activation.

Our study provides a more comprehensive understanding of the molecular subtypes of nAmb neurons than previously known. Two studies previously used single-cell RNA-seq to molecularly classify subtypes of nAmb neurons. For instance, Coverdell et al. (2022), identified three subtypes from 145 *Chat*-expressing nAmb neurons of adult mice ^36^. Another study, Veerakumar et al. (2023), classified three subtypes of nAmb neurons from 203 heart- or larynx-projecting nAmb neurons of neonatal mice ^37^. However, each of these previous studies contained far fewer neurons than the nearly 1,200 found bilaterally in adult mouse nAmb ^50^ and so may have underestimated the diversity of nAmb neurons. Our present study identified eight molecular subtypes of nAmb neurons and one molecular subtype of intermediate-zone cardiovagal neurons among 1,245 Chat-expressing cells, nearly doubling the number of previously recognized nAmb neuron subtypes. However, our results do not exclude the presence of rare nAmb neuron subtypes. Moreover, it remains unclear whether the composition of subtypes differs between the left and right hemispheres of the nAmb or varies with sex, particularly given that the right vagus predominantly controls heart rate more strongly than the left, e.g., ^65^.

Our present study provides a finer resolution of nAmb neuron subtypes than previously known. For instance, a previous study by our group identified three subtypes of nAmb neurons ^5^, which our integrated analysis now resolves as nine subtypes. One subtype identified in our previous dataset, Adcyap1^nAmb^ neurons, matches to multiple subtypes in the combined dataset. Another previously defined subtype, Vipr2^nAmb^ neurons, divides into two subtypes, n5 (Uts2^nAmb^) and n6 (Stk32a^nAmb^), in the present study. Since Vipr2^nAmb^ neurons innervate the pharynx and larynx, future studies could investigate whether the Uts2^nAmb^ and Stk32a^nAmb^ subtypes separately innervate these two upper airway regions. Interestingly, *Uts2* and *Uts2b* expression mark vocal control regions in birds ^66^ and are enriched in the n5/Uts2^nAmb^ cluster of mouse nAmb neurons in the current study, raising the possibility that this subtype innervates the larynx to control vocalization.

Performing label transfer method to our dataset with that of another previous study also sheds light on the functional identities of our nAmb neuron subtypes. Veerakumar and colleagues identified three functional subtypes of nAmb neurons: cardiopulmonary nAmb neurons (ACP), which cause bradycardia and bronchoconstriction; cardiovascular nAmb neurons (ACV), which cause bradycardia without bronchoconstriction; and larynx-projecting nAmb neurons ^37^. Marker gene expression between the current study and Veerakumar *et al.* suggests that clusters n03.s01 (Npy2r^nAmb^ neurons) and n03.s02 of the current study may correspond to ACV neurons (*Bche+*) and ACP (*Calb1+*) neurons, respectively, of the Veerakumar *et al.* study. However, differences in gene expression between the datasets, potentially related to age differences in mice between studies, limit our ability to compare nAmb neuron subtypes between the two studies. Also, while Veerakumar *et al.* did not detect a significant increase in Fos immunofluorescence in ACV neurons after nasal immersion in anesthetized mice, indicating a lack of involvement of ACV neurons in the diving reflex. In contrast, our results show that *Fos* RNA expression increases in Npy2r^nAmb^ neurons in response to voluntary underwater diving. If Npy2r^nAmb^ neurons are ACV neurons, as indicated by our gene expression analysis, then it is possible Npy2r^nAmb^ neurons are activated differently by nasal immersion in the anesthetized state than by voluntary underwater diving.

The activation of Npy2r^nAmb^ neurons during underwater diving is consistent with a well-characterized trigemino-cardiac reflex circuit. Nasal sensory input from the anterior ethmoidal nerve engages neurons in the medullary dorsal horn which then relay to the rostral ventrolateral medulla (RVLM), where cardiovagal preganglionic neurons lie within the external formation of the nucleus ambiguus ^6,26,31,67,68^. Activation of trigeminal afferents evokes a polysynaptic excitatory glutamatergic pathway to CVNs of nAmb ^69,70^, confirming that these neurons serve as direct functional targets of trigemino-cardiac signaling. Together, anatomical tracing, physiological stimulation, and lesion studies point to a short, robust reflex circuit with both direct and indirect pathways from trigeminal sensory neurons to cardiac efferent neurons. These mechanisms provide a framework for understanding how somatosensory stimuli engage parasympathetic output and may help inform therapeutic use of the diving response.

Finally, the molecular profile of Npy2r⁺ nAmb neurons raises hypotheses about signaling pathways through which the vagus nerve controls heart rate. Our analysis identified several genes enriched in this cluster that mediates vagal control of heart rate. For instance, *Prkg1,* which encodes cGMP-dependent protein kinase I, acts as a key effector in nitric oxide (NO) signaling and has been implicated in modulating cardiac contractility and autonomic control of heart rate via NO-cGMP pathways ^71^. Similarly, *Adcy8*, a calcium-sensitive adenylyl cyclase, has been shown to affect heart rate and heart rate variability when overexpressed ^72^. *Ank2*, encoding ankyrin-B, plays a crucial role in targeting cardiac ion channels, transporters, and signaling proteins to the membrane; loss-of-function mutations in *Ank2* can lead to a spectrum of arrhythmias, including sinus node dysfunction, atrial fibrillation, and ventricular arrhythmias, collectively known as “ankyrin-B syndrome” ^73^. Additionally, *Dmd*, which encodes dystrophin, is critical for cardiac integrity, as its deficiency underlies Duchenne cardiomyopathy, characterized by fibrosis, arrhythmias, and heart failure ^74^. Future studies using conditional knockouts could investigate whether these genes control heart rate through their action in Npy2r^nAmb^ neurons.

## Acknowledgements

We gratefully acknowledge Patrice G. Guyenet, Ruth Stornetta, and Steven J. Swoap for thoughtful discussion and feedback on the experimental design; Bradford B. Lowell for the Chat-Flp mouse line; Stephen D. Liberles for the Npy2r-Cre mouse line; Patrice G. Guyenet and Hui Zong for co-acquisition of pilot funding; and Moon Snyder, Natalie Schiavone, and Virginia Owen Trinkle for technical support. We also acknowledge the valuable assistance of these UVA core facilities: the Robert M. Berne Cardiovascular Research Center Histology Core; the Biology Department Genomics Core; the Genome Analysis and Technology Core (RRID:SCR_018883); and the Flow Cytometry Core Facility.

## Funding statement

Funding was provided by a University of Virginia 3 Cavaliers award to J.N.C., Patrice G. Guyenet, and Hui Zong; NIH R01 HL148004 to SBGA; NIH T32 GM007055 and NIH F31 HL158187 to TCC; and a Pathway to Stop Diabetes Initiator Award 1-18-INI-14 and NIH R01 HL153916 to JNC.

## Data availability statement

The raw and processed single-nuclei RNA-seq data and metadata reported in this paper are available at GEO accession number ######. A user-friendly interface for visualizing and exploring the single-nuclei RNA-seq data is available at the Broad Institute Single Cell Portal at ######. The code used for processing, clustering, and visualizing the clustered single-nuclei RNA-seq data is available through Zenodo at ######. Any additional information required to reanalyze the data reported in this paper is available from the lead contact upon reasonable request.

## Conflict of interest disclosure

The authors declare no conflict of interest

## Ethics approval statement

All animal care and experimental procedures were approved in advance by the University of Virginia Institutional Animal Care and Use Committee under animal use protocols #4232 and #4354.

## Patient consent statement

not applicable

## Permission to reproduce material from other sources

not applicable

## Clinical trial registration

not applicable

## AUTHOR CONTRIBUTIONS

MJ, MKM, TCC, VAG, SBGA, and JNC designed the experiments. MJ, TCC, and JNC generated the single-nuclei transcriptomics data and MJ analyzed it. MJ, MKM, TCC, SBGA and MEC performed RNA FISH studies. MEC performed surgeries and chromogenic staining/tissue clearing for the anterograde tracing studies. MJ, AD, MKM, and MEC performed immunofluorescence. YBW and CRB injected the retrograde tracer into the cardiac fat pads. MJ, TCC, and MKM validated the mouse lines. MJ, MKM, MEC, AD, KC, and AS performed electrocardiography. VAG developed the diving reflex assay, DSS performed post-diving histology for ISH, and MEC generated and analyzed diving reflex data. MJ, TCC, MKM, VAG, SBGA, and JNC prepared the figures. MJ, MKM, TCC, SBGA, and JNC wrote the manuscript, with input from all the authors.

## DECLARATION OF INTERESTS

The authors declare no competing interests.

## METHODS

### Experimental Model and Subject Details

The single-cell RNA-seq experiments used 28 - 35 week old male and female B6;129S6-*Chat^tm2(cre)Lowl^*/J mice (“Chat-Cre”; (Jackson Laboratories, JAX, strain # 028861) ^38^, some of which had been crossed to a nuclear reporter mouse line B6.Cg-*Gt(ROSA)26Sortm1(CAG-HIST1H2BJ/mCherry,-EGFP/Rpl10a)Evdr*/J (“H2b-TRAP”; JAX strain #029789) ^75^. The fluorescent *in situ* hybridization experiments used C57BL/6J mice from the Jackson Laboratory (JAX, strain # 000664). The following mouse lines were used for chemogenetic studies: *Npy2rtm1.1(cre)Lbrl*/RcngJ (“Npy2r-Cre”; gift of Stephen Liberles, Harvard Medical School, Howard Hughes Medical Institute) ^44^; and Chat^em1(flp)Lowl^ (“Chat-Flp”; gift of Bradford Lowell, Beth Israel Deaconess Medical Center, Harvard Medical School) ^36^; R26^ds-HTB^ mice (gift of Martyn Goulding, Salk Institute for Biological Studies). The anterograde tracing experiments used Chat-Cre, Npy2r-Cre, and C57BL/6J mice. Unless otherwise specified, all experiments used adult mice with approximately equal numbers of male and female mice. Mice were housed at 22-24 °C with a 12 hour light:12 hour dark cycle and unlimited access to standard mouse chow and water.

### Fluorescently Labeling Nucleus Ambiguus Neurons

To fluorescently label the nAmb *Chat*+ neurons for single-nuclei transcriptomics, two approaches were used. One approach crossed Chat-Cre mice with the H2b-TRAP transgenic mouse line, which Cre-dependently labels cells with mCherry-tagged histone 2B protein (H2b-mCherry) and an eGFP-tagged ribosomal L10 protein. Preliminary transcriptomic analysis failed to detect *Chat* expression in many Chat-Cre;H2b-TRAP-labeled cells (Figure 1A, far right panel), consistent with developmental Chat-Cre activity in brain regions around the nAmb. Therefore, a second labeling approach was used to enrich for nAmb *Chat*+ neurons: an AAV which Cre-dependently expresses H2b-mCherry, AAV9-Syn1-FLEX-H2b-mCherry (Vigene Biosciences, 3.94 × 10^13^ GC/mL), injected into the right ventrolateral medulla of eight male and female Chat-Cre mice, four weeks prior to tissue collection (see “Virus Injections” section for details on surgery).

### Single-Nuclei RNA-Sequencing

Tissue samples of nAmb were collected after rapid decapitation of the mice to avoid stress- and anesthesia-related changes in nuclear mRNA. Brains were immediately extracted, chilled in slush, and sectioned coronally at 1 mm intervals through the nAmb’s full rostral-caudal extent (Bregma −6.5 mm to −8.0 mm). The brain sections were immersed in ice-cold RNAprotect (Qiagen, catalog # 76106) to preserve RNA during microdissection and storage. After at least 30 min in RNAprotect, the nAmb was visualized under a fluorescence stereomicroscope (Zeiss Discovery V8), dissected, and stored in RNAprotect overnight at 4 °C. On the next day, nAmb tissue was homogenized and purified by density-gradient centrifugation into a single-nuclei suspension as previously described ^76,77^. The single-nuclei suspension was sorted by FACS using 80 nm nozzle to collect H2b-mCherry+ nuclei in a 2 mL microcentrifuge tube. The sample was then processed using 10X Genomics Chromium Next-GEM Single Cell 3’ cDNA Kit v3.1 according to the manufacturer’s instructions. 10X cDNA libraries were sequenced on an Illumina Next-Seq 550 with high-output, 75-cycle v2.5 kits. Sequencing reads were demultiplexed by bcl2fastq2 v2.20.0 (Illumina) and aligned to the mouse genome mm10-2020-A with 10X Genomics Cell Ranger software pipeline 5.0.0 using the include-intron function (Supplemental Figure 1C).

Gene expression matrices from the 10X Cell Ranger pipeline were input to Seurat v4.3.0 in R (version 4.1.2) for data analysis. Genes detected in fewer than three cells, cells with fewer than 200 genes, and cells with more than 2% of reads mapping to mitochondrial genes (%mt) were filtered out. Pass-filter data were then merged across batches and integrated to correct for technical differences including batch effects ^78^. The following steps were then performed: log-normalization of the data; selection of the 2,000 highest-variance genes using Seurat’s FindVariableFeatures() function; batch integration using the IntegrateData() function in Seurat; scaling of gene expression values; Principal Component Analysis (PCA) for linear reduction of the dimensionality of data; cell clustering using Louvain algorithm, based on Euclidean distance in the PCA space comprising the first 13 PCs and with a resolution of 1; and non-linear dimensionality reduction by Uniform Manifold Approximation and Projection (UMAP) for visualization in two dimensions ^79^.

To enrich our analysis with nAmb neurons, we included neuron clusters expressing *Slc18a3, Isl1*, and *Chat* and excluded neuron clusters expressing *Slc32a1* and *Slc17a6* (Figure 1A). After removing non-nAmb neurons, the remaining neurons were re-clustered, including the steps of feature selection, PCA, clustering with the top 17 PCs and resolution setting of 0.25. Integration of all nAmb neurons was performed again with 12 PCs at 0.25 resolution. Cluster relatedness in PCA space was illustrated with dendrograms using Seurat’s BuildClusterTree() function.

Differentially expressed genes in each cluster were identified using Seurat’s FindAllMarkers() function with the Wilcoxon Rank Sum test and p values Bonferroni-adjusted for multiple comparisons. The results were filtered to only include positive marker genes expressed in a minimum fraction of 0.25 cells with log2-fold change threshold of 0.25. Candidate marker genes were then manually selected based on their expression specificity in each cluster.

### RNA Fluorescence In Situ Hybridization (FISH)

RNA FISH experiments were performed on brain tissue from mice, some of which received an intraperitoneal injection of 2% Fluorogold (Fluorochrome) to label peripherally projecting neurons, including all nAmb vagal efferent neurons, a minimum of 5 days prior to euthanasia. Mice were terminally anesthetized with ketamine (20 mg/kg) and xylazine (2 mg/kg) diluted in sterile saline, followed by transcardial perfusion with 0.9% saline plus heparin and 4% paraformaldehyde (Thomas Scientific). Brains were extracted and post-fixed in 4% paraformaldehyde for 24 hr at 4 °C. Following fixation, brains were sectioned coronally at 30 - 35 um thickness on a vibratome (VT-1000S, Leica). The day before FISH, the sections were rinsed in PBS and then mounted on precleaned Superfrost Plus microscope slides (Fisher, catalog # 12-550-15) and left to dry overnight. An ImmEdge Hydrophobic Barrier Pen was used to draw a barrier around the sections. The sections were then incubated in Protease IV in a HybEZ II Oven for 30 min at 40°C, followed by incubation with target probes (*Chat*, *Phox2b*, *Adcyap1*, *Npy2r*, *Phox2b, Cre,* and *Flp*) for 2 hr at 40 °C. Slides were then treated with AMP 1-3, HRP-C1, HRP-C2, HRP-C3, and HRP Blocker for 15-30 min at 40 °C, as previously described ^80^. FITC, Cy3, and Cy5 (Perkin Elmer) fluorophores were used for probe visualization. Fluorogold was imaged based on its native fluorescence. Images were taken using a confocal microscope (Zeiss LSM 800). The three levels of the nucleus ambiguus correspond to the following bregma levels: rostral or compact nAmb: −6.47 mm through −6.75 mm from bregma; intermediate or semi-compact nAmb: −6.83 mm through −7.10 mm from bregma; caudal or loose nAmb: −7.19 mm through −7.46 mm from bregma.

### Virus Injections

Mice were generally anesthetized with ketamine (20 mg/kg) and xylazine (2 mg/kg), positioned in a stereotaxic apparatus (Kopf), and locally anesthetized with long-lasting bupivicaine. A pulled glass micropipette was used with a Nanoject III system to stereotaxically inject one of the following viral vectors per mouse: AAV9-CAG-FLEX-tdTomato (“AAV9-FLEX-tdTomato”), AddGene plasmid # 28306, RRID:Addgene_28306, a gift from Edward Boyden; AAV9-CAG-FLEX-PLAP virus (“AAV9-FLEX-PLAP”), a gift from Steve Liberles and Sara Prescott; AAV9-Syn1-DIO-H2b-mCherry (“AAV9-DIO-H2b-mCherry”), Vigene Biosciences, 3.94 × 10^13^ GC/m; or AAV1/2-CreON/FlpON-hM3Dq-HA, Viral Vector Facility, University of Zurich and ETH Zurich, catalog # vhW34-1. To span the rostrocaudal extent of the nAmb, multiple injections were made at the following coordinates relative to the calamus scriptorus: anterior/posterior −2.1, −2.4, −2.7 mm, lateral/medial +/− 1.3 mm, and dorsal/ventral −5.8 mm, from lambda; and anterior/posterior - 0.1 mm, lateral/medial +/− 1.3 mm, and dorsal/ventral −0.1, −1.3 mm. AAV9-FLEX-tdTomato and AAV9-DIO-H2b-mCherry virus was injected unilaterally into the ventrolateral medulla, 50 nL per injection, 3 injections on the right side per mouse. AAV9-FLEX-PLAP and AAV1/2-CreOn/FlpOn-hM3Dq-HA were injected bilaterally, 50 nL per injection, 6 injections total per mouse. This injection strategy was designed to fully cover the nAmb, avoid the DMV, and restrict viral infection to only nAmb neurons. The pipette was removed 5 minutes after each injection, followed by wound closure with sutures or surgical wound glue (Vetbond). Meloxicam SR (5 mg/kg; sustained release, SR) was injected subcutaneously for post-operative analgesia.

### Tissue Histology and Immunofluorescence

Three to four weeks after AAV9-FLEX-tdTomato or AAV9-FLEX-PLAP virus injection, mice were deeply anesthetized with ketamine (20 mg/kg) and xylazine (2 mg/kg) diluted in sterile saline, then transcardially perfused with 0.9% saline plus heparin followed by 4% paraformaldehyde (PFA) (Thomas Scientific; CAS#30525-89-4) or 10% Neutral Buffered Formalin solution; NBF (Sigma-Aldrich HT501128-4L). Brains and hearts were harvested and shaken in the same fixative overnight at room temperature. Brains were either sectioned on vibratome at 35 um or cryoprotected in 30% sucrose on a shaker at 4 °C overnight and then sectioned on freezing microtome at 35 um. Hearts were cryoprotected in 30% sucrose solution in 4 °C on a shaker for 2 days. Hearts were then cut with a 10 blade (Fine Science Tools, Catalog #10010-00) along the dorsal ventral axis and were then embedded in optimal cutting temperature (OCT) compound and sectioned on cryostat (Thermo Scientific CryoStar NX50) at 10 um intervals. Slide-mounted cardiac sections were then kept at –80 °C for long term storage.

For heart immunofluorescence, slides were removed from –80 °C and placed flat on a tray in a fume hood to dry at room temperature. A hydrophobic barrier was drawn to contain solutions on the slide, and the slides were left to air dry for 10 min. The slides were then placed in the pre-chilled Coplin jar containing acetone in –20 °C for 10 min. After 10 min, acetone was removed, and the slides dried in the fume hood for 10 min. A 0.1% Sudan Black solution dissolved in 70% ethanol was added for 20 min to reduce autofluorescence. Tissue was then washed three times for 5 min with PBS. 0.1% PBST was prepared by dissolving 100 uL of Triton-X100 (Sigma Life Sciences, catalog # 9036-19-5) in 100 ml 1X PBS (Fisher, catalog # 70-013-032). Chat-Cre and Npy2r-Cre heart tissues were incubated in a blocking buffer for an hour on a shaker at room temperature. The blocking buffer consisted of 5% donkey serum (Millipore Sigma, catalog # S30-100ML) in 0.1% PBST. For anterograde tracing with AAV-tdTomato, the tissue was incubated overnight at 4 °C on a shaker with primary antibodies against PGP9.5 (rabbit, Abcam, catalog #EPR4118) and tdTomato (goat, Arigo Biolaboratories, catalog #ARG55724), both diluted 1:500 in 5% Donkey blocking buffer. Similarly, for anterograde tracing with AAV-PLAP, the tissue was incubated with anti-PLAP (rabbit, Abcam, catalog #AB133602) and anti-ChAT (goat, Sigma Aldrich, catalog #AB144P), diluted 1:500 and 1:100 respectively, in 5% donkey blocking buffer.

For brain immunohistochemistry, tissue was rinsed in PBS three times for 5 minutes on a shaker at room temperature, followed by incubation in 5% blocking buffer (5% normal donkey serum + 0.1% PBST) for 1 hour at room temperature on the shaker. Brain sections were then incubated in primary antibody against ChAT (from goat; Sigma Aldrich, catalog # AB144P; 1:100 dilution), PLAP (from rabbit, catalog # AB133602; 1:500 dilution) and anti-HA (from chicken, Catalog # ET-HA100; dilution 1:500).

After primary antibody incubation, heart and brain sections were then rinsed three times for 5 minutes in 0.1% PBST solution. Species-specific donkey secondary antibodies conjugated to Alexa Fluor 488, 568 or 647 were obtained from Abcam or Invitrogen and heart and brain sections were incubated in secondary antibodies (dilution 1:1000) for 2-4 hours on a shaker at room temperature. After three 5 minutes in PBS washes, the tissue was mounted on Superfrost Plus precleaned slides (Fisher; catalog # 12-550-15) followed by cover-slipping with Prolong Gold antifade mounting media (Cell Signaling Technology, catalog # 9071S) and sealed with nail polish.

### Retrograde Labeling of Heart-Projecting Neurons

Mice were deeply anesthetized using isoflurane (2.5%) and placed in a supine position on a heating pad. A thoracotomy was performed by making a small incision between the second and third ribs of the right rib cages to expose the heart. A 40 uL volume of Fluorogold (1%; Fluorochrome) was injected into the pericardiac fat pad near the posterior right of atrioventricular junction where cardiovagal nerve endings terminate. A small cotton swab was used to absorb any excess tracer to minimize non-specific labeling. The incision was closed using suture and animals were returned to their home cage for recovery after surgery and monitored for 3 days. Carprofen (10 mg/kg) and buprenorphine (0.1 mg/kg) were administered immediately after surgery and one day post-surgery.

After week animals were perfused with ketamine (20 mg/kg) and xylazine (2 mg/kg) diluted in sterile saline, followed by transcardial perfusion with 0.9% saline and heparin then 10% neutral buffered formalin solution (“NBF”, Sigma-Aldrich HT501128-4L). The brains were collected and post-fixed for 24 hr at room temperature in 10% NBF. Following fixation, brains were transferred to 30% sucrose solution for 48 hr at 4 °C on a shaker and sectioned coronally at 30 or 35 um thickness on a microtome (SM2010R, Leica Biosystems) for immunofluorescence and RNA FISH as described in the corresponding sections above.

### Placental Alkaline Phosphatase Staining

Following 3-4 weeks after AAV9-FLEX-PLAP virus injection ^59^, mice were terminally anesthetized with ketamine (20 mg/kg) and xylazine (2 mg/kg) diluted in sterile saline, followed by transcardial perfusion with 0.9% saline plus heparin then 10% neutral buffered formalin solution (“NBF”; Sigma-Aldrich HT501128-4L). The brains, esophagus, trachea, larynx, pharynx and lungs were collected and post-fixed for 24 hr at 4 °C in 10% NBF. Following fixation, brains were transferred to 30% sucrose solution for 48 hr at 4 °C on a shaker and sectioned coronally at 30 um or 35 um thickness on a microtome (SM2010R, Leica Biosystems). A single series of brain sections per mouse was used in histological studies to confirm injection site and PLAP expression. All mice with little to no reporter expression in the nAmb were determined to be surgical “misses’’ and so excluded from anatomical analyses.

En bloc tissue samples of the esophagus and upper airways were washed three times for 1 hour at room temperature in PBS, followed by incubation in alkaline phosphatase (AP) buffer (0.1 M Tris HCl pH 9.5, 0.1 M NaCl, 50 mM MgCl2, 0.1% Tween 20, 5 mM tetramisole-HCl) for two hours at 70 °C. Afterward, the samples were equilibrated to room temperature and then washed twice in AP buffer. AP activity was visualized with NCT/BCIP solution (ThermoFisher Scientific 34042) and stained samples were rinsed in AP buffer for 15 min, post-fixed in 10% NBF for 1 hr, and washed in PBS. Samples were then dehydrated through a series of ethanol washes (15% - 100%) and cleared using a 1:2 mixture of benzyl alcohol (Sigma-Aldrich 402834-500ML) and benzyl benzoate (Sigma-Aldrich B6630-1L). Whole mount images were taken using a brightfield stereomicroscope (Leica M205 FCA Stereomicroscope with color camera) and brain sections were imaged using fluorescence microscope (Echo Revolve).

### Chemogenetic Experiments during Non-Invasive Electrocardiography

AAV1/2-CreON/FlpON-hM3dq-HA was injected into the bilateral nAmb of Npy2r-Cre;Chat-Flp;R26ds-HTB mice as described above. The mice were allowed to recover for 6 to 8 weeks before further experimentation. A non-invasive electrocardiography system (ECGenie) was then used to measure heart rate. All experiments were performed at the same time of day to avoid circadian differences in heart rate. Mice were acclimated to the ECGenie recording towers for three days prior to data collection. Deschloroclozapine (Tocris, Cat #7193) was prepared by dissolving it in DMSO, then diluting with 0.9% sterile saline to obtain a 2% DMSO solution of DCZ. DCZ-vehicle was prepared by dissolving 2% DMSO in 0.9% sterile saline. After the acclimation period, mice were placed into the ECGenie recording towers for a 10 min baseline recording of heart rate using Corvita data acquisition software (Mouse Specifics, Framingham, MA, version 2.09). Afterwards, DCZ-vehicle was administered by intraperitoneal injection, and heart rate was recorded for 20 min. DCZ was then injected (5 mg/kg; i.p.), and heart rate was recorded for 60 min. Next, volume-matched injections of either vehicle (0.9% sterile saline) or methyl atropine bromide (MA; 1 mg/kg, dissolved in 0.9% sterile saline) was administered, and heart rate was recorded for 30 minutes. Each mouse underwent at least one trial in which MA was injected after DCZ and at least one trial in which MA-vehicle was injected after DCZ. Data was analyzed using EzCG Analysis Software (Mouse Specifics, Framingham, MA, version 17). For all heart rate recordings, 3 - 5 seconds of ECG data was collected every three minutes. ECG signals were then visually inspected for missed or ectopic beat detection. Pieces of data with noisy signal or incorrect beat detection were removed before calculating heart rate, and if needed, the signal was inverted.

Heart rate values were averaged per animal for each condition. Comparisons between baseline, vehicle, and DCZ conditions were performed using one-way repeated-measures ANOVA, followed by Tukey’s honestly significant difference test for post-hoc pairwise comparisons using the emmeans package in R (version 4.1.2), which computes exact Tukey-adjusted p-values. For the comparisons of methyl-atropine (MA) versus MA-vehicle conditions, paired t-tests were performed. A p-value < 0.05 was considered statistically significant.

To validate expression of hM3Dq, mice were euthanized, transcardially perfused, and fixed as described above after completion of the ECG studies. Brains were fixed overnight, transferred to 30% sucrose solution for 48 hr at 4°C on a shaker, and sectioned coronally at 30 or 35 um thickness on a microtome (SM2010R, Leica Biosystems). Sections were rinsed 3 times for 5 minutes in PBS, then incubated in blocking buffer (5% normal donkey serum + 0.1% PBST) for 1 hour on a shaker at room temperature. After blocking, sections were incubated overnight at 4 °C on a shaker in primary antibody solution containing goat anti-Chat (Primary antibody at 1:100 dilution in blocking buffer, Sigma Aldrich catalog # AB144P, lot # 3491643) and chicken anti-HA antibody (1:500 dilution, Aves Lab, lot # HA7757994, catalog # ET-HA 100). The following day, sections were rinsed 3 times for 5 minutes in 0.1 % PBST solution (100 uL Triton-X dissolved in 100 mL sterile PBS) on a shaker, then incubated in secondary antibody for 2-4 hours at room temperature. Donkey anti-chicken Alexa Fluor 647 (Jackson ImmunoResearch, reference # 703-585-155, lot # 147524; 1:1000 in 0.1% PBST) and donkey anti-goat, Fluorophore 488 (Invitrogen, Ref# A11055, Lot#2747580; 1:1000 in 0.1% PBST). Sections were then rinsed in PBS 3 times for 5 minutes on a shaker at room temperature. After rinsing, sections were mounted onto slides, allowed to dry, and cover-slipped using Vectashield Antifade Mounting Medium with DAPI (Cat# H-1200-10), then sealed with nail polish. Slides were imaged using the confocal microscope. Mice were excluded from functional analysis if their nAmb lacked hM3Dq-HA immunofluorescence, or their DMV contained more than 5 hM3Dq-HA immunofluorescent neurons.

### Voluntary Dive Training

Npy2r-Cre;Chat-Flp;dsHTB and C57Bl/6J mice were trained to voluntarily dive underwater according to a protocol adapted from a previous publication ^81^. A diving pool was built from a 69 cm long x 38 cm wide × 10 cm high plastic tub. A channel was created in the center of the pool by affixing two 69 cm long x 0.32 cm wide × 10 cm high plexiglass strips with 2.5 cm perpendicular ledges 6 cm from the bottom, one third and two thirds of the way across the length of the pool. These served as rails along which water-level barriers were placed to extend the length of the underwater dive. Squares of 10 cm x 0.6 cm × 10 cm plexiglass were used to construct walls around the starting end of the pool. To set up the diving enclosure for training, the two plexiglass strips were taped in place onto either end of the container and water was added to the tank to the desired level. Water temperatures were kept between 33 °C and 37 °C to prevent hypothermia and minimize stress. A raised platform was placed at the opposite end for the mouse to climb out of the water (escape platform). Small circles were drawn every 15 cm on the inside walls of the two plexiglass strips to help guide the mice while underwater.

Additionally, a large “X” was drawn on the underwater portion of the platform for the same purpose. The diving enclosure was set up with the platform and thermometer and filled with water before every dive training session; and drained and cleaned after every session. A heat lamp was set up approximately 30 cm above the recovery cages to provide warmth after dive completion. At the start of each swim or dive, the mouse was taken from its home cage, placed at the starting end of the pool, allowed to swim across to the escape platform, and then removed from the pool and placed into its home cage under the warming lamp.

A 5-day progressive protocol for dive training mice was performed as follows. On Day 1 of dive training, mice were allowed 2 min to explore the pool with 1 cm of water. If a mouse touched the escape platform with all 4 paws within 2 min, they were immediately removed from the pool. If they failed to reach the escape platform, they were removed after 2 min. This was repeated 7 times. On Day 2 of dive training, mice were allowed to explore with the same criteria as in Day 1, for two trials. Then, water was added up to 3 cm and mice were allowed to wade through the water, for two trials. Water was added up to 5 cm, so the mice were unable to touch the bottom of the enclosure while swimming, for 7 trials. On Day 3, mice underwent a series of underwater dive trials of increasing distances: one trial of a 1 cm long dive; two trials of a 3 cm long dive; four trials of a 5 cm long dive; and six trials of a 6 cm long dive. On Day 4, mice underwent one trial each of 1 cm, 3 cm and 5 cm long dives, two trials of a 1cm long dive, and 5 trials of a 10 cm long dive. On Day 5, mice did one trial each of 1 cm, 3 cm and 5 cm long dives, two trials each of a 1 cm and a 10 cm long dive, and five trials of a 20 cm long dive. Dive repetitions were increased or decreased depending on each mouse’s learning speed.

### Telemetric Electrocardiography

After dive training completion, mice underwent surgical implantation of electrocardiography (EKG) telemeters (Data Science International, DSI; ETA-F10) for wirelessly recording heart rate at rest and during diving. Each mouse was anesthetized with isoflurane and placed in dorsal recumbency on the auto-heated surgical table. The skin on the chest and upper abdomen was shaved, cleaned with alcohol and iodine, and 0.1mL of Nocita (long-lasting Bupivacaine, 13.3 mg/mL) was injected at the midline of the sternum, with the limbs loosely taped down. A 2 cm shallow, vertical inline incision was made along the sternum and, using blunt-tip hemostatic forceps, a subcutaneous pocket starting at the left caudal edge of the incision was created down the lateral flank. Once the pocket size was increased to ensure adequate fit for the probe, the pocket and the probe were rinsed with sterile saline. The device ID number was noted alongside the mouse ID and the probe was inserted into the pocket with the leads oriented cranially (red lead on the left and white lead on the right). The leads were trimmed to the appropriate length to allow for placement in desired locations. The red lead was routed subcutaneously by blunt dissection to the left flank inferior to the underarm and secured with a 6.0 silk suture to the muscle. The white lead was routed subcutaneously across the torso to the upper right pectoralis major and secured with a 6.0 silk suture. Secure attachment of the probe was confirmed by slightly tugging on the leads. The incision was closed with 3 mouse wound clips and antibiotic ointment was applied. The wound clips were reapplied if dislodged by the mouse. Wound clips were subsequently removed 7-10 days later.

### Diving Reflex Assay

For diving-induced *Fos* experiments, a total of 13 mice (7 females, 6 males) were trained according to the above protocol. Before training, mice were administered Fluorogold systemically (FG; i.p. as described above) to label nAmb neurons. After 5-7 days of training, mice were randomly assigned into groups (voluntary dive or no dive). For voluntary diving, 7 mice (4 females, 3 males) were introduced into the diving chamber filled with water and allowed to dive voluntarily to reach the platform. Upon reaching the platform, mice were given 10-15 sec rest before being returned to the opposite end of the apparatus to dive again. For 20-30 min mice completed 20-40 individual dives before being placed in a warmed recovery chamber for 20 min to allow time for *Fos* mRNA expression. As a control (‘no dive’), mice were placed in the diving enclosure filled with 1 cm of water and allowed to explore for 30 min before being placed in a warmed recovery chamber for 20 min before perfusion.

Mice were deeply anesthetized with a mixture of ketamine (100 mg/kg) and dexmedetomidine (0.2 mg/kg) given i.p. and perfused transcardially with 4% paraformaldehyde, pH 7.4 in 100 mM phosphate buffers. Brains were removed and post-fixed in the same fixative for 12-24 hr at 4 °C. Brains were sectioned (30-50 µm) on a vibratome (VT-1000S, Leica Biosystems, Deer Park, IL, USA), and sections were stored in cryoprotectant (30% ethylene glycol (v/v), 20% glycerol (v/v), 50% 100mM phosphate buffer, pH 7.4) at −20°C. Multiplex fluorescent in situ hybridization was performed using RNAscope (V1 kit, Advanced Cell Diagnostics, Newark, CA, USA). Probes for *Npy2r*, *Chat*, and *Fos*, and native fluorescence was used to identify Fluorogold+ cells. Serial 1-in-3 sections were washed in RNase(ribonuclease)-free phosphate buffered saline, mounted on charged slides, dried overnight, and processed according to the manufacturer’s instructions.

Immediately following the RNAscope procedure, sections were rinsed and then incubated in blocking solution for 10 min followed by incubation in primary at room temperature for 60 min, rinsed and incubated in secondary antibodies for 30 min, rinsed and then dried overnight before cover slipping. Slides were cover slipped with Prolong Gold antifade mounting media with DAPI (Thermo Fisher Scientific, catalog # P3693).

All nAmb neurons on one side of the brain between the spinal decussation and facial nucleus were counted. Only cells that contained FG fluorescence and stained for *Chat* RNA were included. Presence or absence of *Npy2r* and *Fos* RNA was determined for each FG+/*Chat*+ nAmb neuron. A cell was considered *Fos*+ if it contained >5 bright *Fos* puncta.

## Notes

### Competing Interest Statement

The authors have declared no competing interest.

### Summary of Updates

The revised manuscript now includes additional experiments involving retrograde tracing of heart-projecting neurons, intersectional chemogenetic activation of a molecular subtype of heart-projecting neuron, and anterograde tracing from the same neuron subtype to the upper airways and esophagus, as well as relevant control experiments and experimental validation of methods.

## REFERENCES

1 Weber, E. a. E. H. Experiences qui prouvent que les nerfs vague, stimulés par l’appariel de rotation galvano-magnetique, peuvent retarder et même arrêter le movement du coeur. Arch. gen. Med. (1846).

2 Fye, W. B. Profiles in cardiology. Ernst, Wilhelm, and Eduard Weber. Clin Cardiol 23, 709–710, doi:10.1002/clc.4960230915 (2000).

3 Dampney, R. A. Central neural control of the cardiovascular system: current perspectives. Adv Physiol Educ 40, 283–296, doi:10.1152/advan.00027.2016 (2016).

4 Ottaviani, M. M. & Macefield, V. G. Structure and Functions of the Vagus Nerve in Mammals. Compr Physiol 12, 3989–4037, doi:10.1002/cphy.c210042 (2022).

5 Coverdell, T. C., Abbott, S. B. G. & Campbell, J. N. Molecular cell types as functional units of the efferent vagus nerve. Semin Cell Dev Biol 156, 210–218, doi:10.1016/j.semcdb.2023.07.007 (2024).

6 Panneton, W. M. & Gan, Q. The Mammalian Diving Response: Inroads to Its Neural Control. Front Neurosci 14, 524, doi:10.3389/fnins.2020.00524 (2020).

7 Goodwyn, E. Dissertatio medica de morteque submersorum investigandis. Department of Medicine, University of Edinburgh, Edinburgh (1786).

8 Vega, J. L. Edmund Goodwyn and the first description of diving bradycardia. Journal of Applied Physiology 123, 275–277, doi:10.1152/japplphysiol.00221.2017 (2017).

9 Butler, P. J. & Jones, D. R. Physiology of diving of birds and mammals. Physiol Rev 77, 837–899, doi:10.1152/physrev.1997.77.3.837 (1997).

10 Panneton, W. M. The mammalian diving response: an enigmatic reflex to preserve life? Physiology (Bethesda) 28, 284–297, doi:10.1152/physiol.00020.2013 (2013).

11 Alboni, P., Alboni, M. & Gianfranchi, L. Diving bradycardia: a mechanism of defence against hypoxic damage. J Cardiovasc Med (Hagerstown) 12, 422–427, doi:10.2459/JCM.0b013e328344bcdc (2011).

12 Geis, G. S. & Wurster, R. D. Horseradish peroxidase localization of cardiac vagal preganglionic somata. Brain research 182, 19–30, doi:10.1016/0006-8993(80)90827-6 (1980).

13 Stuesse, S. L. Origins of cardiac vagal preganglionic fibers: a retrograde transport study. Brain research 236, 15–25 (1982).

14 Standish, A., Enquist, L. W. & Schwaber, J. S. Innervation of the heart and its central medullary origin defined by viral tracing. Science 263, 232–234, doi:10.1126/science.8284675 (1994).

15 Cheng, Z. & Powley, T. L. Nucleus ambiguus projections to cardiac ganglia of rat atria: an anterograde tracing study. The Journal of comparative neurology 424, 588–606 (2000).

16 Cheng, Z., Powley, T. L., Schwaber, J. S. & Doyle, F. J., 3rd. Projections of the dorsal motor nucleus of the vagus to cardiac ganglia of rat atria: an anterograde tracing study. The Journal of comparative neurology 410, 320–341 (1999).

17 Li, Y. F., LaCroix, C. & Freeling, J. Specific subtypes of nicotinic cholinergic receptors involved in sympathetic and parasympathetic cardiovascular responses. Neurosci Lett 462, 20–23, doi:10.1016/j.neulet.2009.06.081 (2009).

18 Levy, M. N. Sympathetic-parasympathetic interactions in the heart. Circ Res 29, 437–445 (1971).

19 Calupca, M. A., Vizzard, M. A. & Parsons, R. L. Origin of pituitary adenylate cyclase-activating polypeptide (PACAP)-immunoreactive fibers innervating guinea pig parasympathetic cardiac ganglia. The Journal of comparative neurology 423, 26–39 (2000).

20 Tompkins, J. D., Ardell, J. L., Hoover, D. B. & Parsons, R. L. Neurally released pituitary adenylate cyclase-activating polypeptide enhances guinea pig intrinsic cardiac neurone excitability. J Physiol 582, 87–93, doi:10.1113/jphysiol.2007.134965 (2007).

21 Feliciano, L. & Henning, R. J. Vagal nerve stimulation releases vasoactive intestinal peptide which significantly increases coronary artery blood flow. Cardiovasc Res 40, 45–55, doi:10.1016/s0008-6363(98)00122-9 (1998).

22 Shanks, J., Pachen, M., Chang, J. W., George, B. & Ramchandra, R. Cardiac vagal nerve activity increases during exercise to enhance coronary blood flow. Circulation research 133, 559–571 (2023).

23 Reed, C. Effects of bilateral vagotomy on blood pressure. American Journal of Physiology-Legacy Content 74, 61–69 (1925).

24 Reed, C. & Layman, J. Effects of bilateral vagotomy on blood pressure and heart rate. American Journal of Physiology-Legacy Content 92, 275–281 (1930).

25 Harvey, R. D. Muscarinic receptor agonists and antagonists: effects on cardiovascular function. Handb Exp Pharmacol, 299–316, doi:10.1007/978-3-642-23274-9_13 (2012).

26 Thomas, M. R. & Calaresu, F. R. Localization and function of medullary sites mediating vagal bradycardia in the cat. The American journal of physiology 226, 1344–1349, doi:10.1152/ajplegacy.1974.226.6.1344 (1974).

27 Chen, H. I. & Chai, C. Y. Integration of the cardiovagal mechanism in the medulla oblongata of the cat. The American journal of physiology 231, 454–461, doi:10.1152/ajplegacy.1976.231.2.454 (1976).

28 McAllen, R. M. & Spyer, K. M. The location of cardiac vagal preganglionic motoneurones in the medulla of the cat. J Physiol 258, 187–204 (1976).

29 Geis, G. S. & Wurster, R. D. Cardiac responses during stimulation of the dorsal motor nucleus and nucleus ambiguus in the cat. Circ Res 46, 606–611, doi:10.1161/01.res.46.5.606 (1980).

30 Geis, G. S., Kozelka, J. W. & Wurster, R. D. Organization and reflex control of vagal cardiomotor neurons. J Auton Nerv Syst 3, 437–450, doi:10.1016/0165-1838(81)90080-1 (1981).

31 Panneton, W. M., Anch, A. M., Panneton, W. M. & Gan, Q. Parasympathetic preganglionic cardiac motoneurons labeled after voluntary diving. Frontiers in Physiology 5, 8 (2014).

32 Lawn, A. M. The localization, in the nucleus ambiguus of the rabbit, of the cells of origin of motor nerve fibers in the glossopharyngeal nerve and various branches of the vagus nerve by means of retrograde degeneration. The Journal of comparative neurology 127, 293–306, doi:10.1002/cne.901270210 (1966).

33 Holstege, G., Graveland, G., Bijker-Biemond, C. & Schuddeboom, I. Location of motoneurons innervating soft palate, pharynx and upper esophagus. Anatomical evidence for a possible swallowing center in the pontine reticular formation. An HRP and autoradiographical tracing study. Brain Behav Evol 23, 47–62, doi:10.1159/000121488 (1983).

34 Fryscak, T., Zenker, W. & Kantner, D. Afferent and efferent innervation of the rat esophagus. A tracing study with horseradish peroxidase and nuclear yellow. Anat Embryol (Berl) 170, 63–70, doi:10.1007/BF00319459 (1984).

35 Bieger, D. & Hopkins, D. A. Viscerotopic representation of the upper alimentary tract in the medulla oblongata in the rat: the nucleus ambiguus. The Journal of comparative neurology 262, 546–562, doi:10.1002/cne.902620408 (1987).

36 Coverdell, T. C., Abraham-Fan, R. J., Wu, C., Abbott, S. B. G. & Campbell, J. N. Genetic encoding of an esophageal motor circuit. Cell Rep 39, 110962, doi:10.1016/j.celrep.2022.110962 (2022).

37 Veerakumar, A., Yung, A. R., Liu, Y. & Krasnow, M. A. Molecularly defined circuits for cardiovascular and cardiopulmonary control. Nature 606, 739–746, doi:10.1038/s41586-022-04760-8 (2022).

38 Rossi, J. et al. Melanocortin-4 receptors expressed by cholinergic neurons regulate energy balance and glucose homeostasis. Cell metabolism 13, 195–204, doi:10.1016/j.cmet.2011.01.010 (2011).

39 Tanaka, I., Ezure, K. & Kondo, M. Distribution of glycine transporter 2 mRNA-containing neurons in relation to glutamic acid decarboxylase mRNA-containing neurons in rat medulla. Neurosci Res 47, 139–151 (2003).

40 Sherman, D., Worrell, J. W., Cui, Y. & Feldman, J. L. Optogenetic perturbation of preBotzinger complex inhibitory neurons modulates respiratory pattern. Nature neuroscience 18, 408–414, doi:10.1038/nn.3938 (2015).

41 Anderson, T. M. et al. A novel excitatory network for the control of breathing. Nature 536, 76–80, doi:10.1038/nature18944 (2016).

42 Herring, N., Lokale, M. N., Danson, E. J., Heaton, D. A. & Paterson, D. J. Neuropeptide Y reduces acetylcholine release and vagal bradycardia via a Y2 receptor-mediated, protein kinase C-dependent pathway. J Mol Cell Cardiol 44, 477–485, doi:10.1016/j.yjmcc.2007.10.001 (2008).

43 Smith-White, M. A., Herzog, H. & Potter, E. K. Role of neuropeptide Y Y(2) receptors in modulation of cardiac parasympathetic neurotransmission. Regulatory peptides 103, 105–111, doi:10.1016/s0167-0115(01)00368-8 (2002).

44 Chang, R. B., Strochlic, D. E., Williams, E. K., Umans, B. D. & Liberles, S. D. Vagal Sensory Neuron Subtypes that Differentially Control Breathing. Cell 161, 622–633, doi:10.1016/j.cell.2015.03.022 (2015).

45 Hao, Y. et al. Integrated analysis of multimodal single-cell data. Cell 184, 3573–3587 e3529, doi:10.1016/j.cell.2021.04.048 (2021).

46 Shuto, Y., Uchida, D., Onda, H. & Arimura, A. Ontogeny of pituitary adenylate cyclase activating polypeptide and its receptor mRNA in the mouse brain. Regulatory peptides 67, 79–83, doi:10.1016/s0167-0115(96)00116-4 (1996).

47 Schmued, L. C. & Fallon, J. H. Fluoro-Gold: a new fluorescent retrograde axonal tracer with numerous unique properties. Brain research 377, 147–154, doi:10.1016/0006-8993(86)91199-6 (1986).

48 Merchenthaler, I. Neurons with access to the general circulation in the central nervous system of the rat: a retrograde tracing study with fluoro-gold. Neuroscience 44, 655–662, doi:10.1016/0306-4522(91)90085-3 (1991).

49 Kalia, M. Brain stem localization of vagal preganglionic neurons. J Auton Nerv Syst 3, 451–481, doi:10.1016/0165-1838(81)90081-3 (1981).

50 Sturrock, R. R. A comparison of age-related changes in neuron number in the dorsal motor nucleus of the vagus and the nucleus ambiguus of the mouse. J Anat 173, 169–176 (1990).

51 Strain, M. M. et al. Dorsal motor vagal neurons can elicit bradycardia and reduce anxiety-like behavior. iScience 27, 109137, doi:10.1016/j.isci.2024.109137 (2024).

52 Henning, R. J. & Sawmiller, D. R. Vasoactive intestinal peptide: cardiovascular effects. Cardiovascular research 49, 27–37 (2001).

53 Weihe, E., Reinecke, M. & Forssmann, W. Distribution of vasoactive intestinal polypeptide-like immunoreactivity in the mammalian heart: interrelation with neurotensin-and substance P-like immunoreactive nerves. Cell and tissue research 236, 527–540 (1984).

54 Grkovic, I., Fernandez, K., McAllen, R. M. & Anderson, C. R. Misidentification of cardiac vagal pre-ganglionic neurons after injections of retrograde tracer into the pericardial space in the rat. Cell Tissue Res 321, 335–340, doi:10.1007/s00441-005-1145-1 (2005).

55 Nosaka, S., Yamamoto, T. & Yasunaga, K. Localization of vagal cardioinhibitory preganglionic neurons with rat brain stem. The Journal of comparative neurology 186, 79–92, doi:10.1002/cne.901860106 (1979).

56 Richardson, R. J., Grkovic, I. & Anderson, C. R. Immunohistochemical analysis of intracardiac ganglia of the rat heart. Cell Tissue Res 314, 337–350, doi:10.1007/s00441-003-0805-2 (2003).

57 Haxhiu, M. A., Jansen, A. S., Cherniack, N. S. & Loewy, A. D. CNS innervation of airway-related parasympathetic preganglionic neurons: a transneuronal labeling study using pseudorabies virus. Brain research 618, 115–134 (1993).

58 McGovern, A. E. & Mazzone, S. B. Characterization of the vagal motor neurons projecting to the Guinea pig airways and esophagus. Front Neurol 1, 153, doi:10.3389/fneur.2010.00153 (2010).

59 Prescott, S. L., Umans, B. D., Williams, E. K., Brust, R. D. & Liberles, S. D. An airway protection program revealed by sweeping genetic control of vagal afferents. Cell 181, 574–589. e514 (2020).

60 Prescott, S. L., Umans, B. D., Williams, E. K., Brust, R. D. & Liberles, S. D. An Airway Protection Program Revealed by Sweeping Genetic Control of Vagal Afferents. Cell 181, 574–589 e514, doi:10.1016/j.cell.2020.03.004 (2020).

61 Izzo, P. N., Deuchars, J. & Spyer, K. M. Localization of cardiac vagal preganglionic motoneurones in the rat: immunocytochemical evidence of synaptic inputs containing 5-hydroxytryptamine. The Journal of comparative neurology 327, 572–583, doi:10.1002/cne.903270408 (1993).

62 Bourane, S. et al. Identification of a spinal circuit for light touch and fine motor control. Cell 160, 503–515, doi:10.1016/j.cell.2015.01.011 (2015).

63 Lee, B. H., Lynn, R. B., Lee, H. S., Miselis, R. R. & Altschuler, S. M. Calcitonin gene-related peptide in nucleus ambiguus motoneurons in rat: viscerotopic organization. The Journal of comparative neurology 320, 531–543, doi:10.1002/cne.903200410 (1992).

64 Nagai, Y. et al. Deschloroclozapine, a potent and selective chemogenetic actuator enables rapid neuronal and behavioral modulations in mice and monkeys. Nature neuroscience 23, 1157–1167, doi:10.1038/s41593-020-0661-3 (2020).

65 Muppidi, S., Gupta, P. K. & Vernino, S. Reversible right vagal neuropathy. Neurology 77, 1577–1579, doi:10.1212/WNL.0b013e318233b3a2 (2011).

66 Bell, Z. W. et al. Urotensin-related gene transcripts mark developmental emergence of the male forebrain vocal control system in songbirds. Sci Rep 9, 816, doi:10.1038/s41598-018-37057-w (2019).

67 Ciriello, J. & Calaresu, F. R. Medullary origin of vagal preganglionic axons to the heart of the cat. J Auton Nerv Syst 5, 9–22, doi:10.1016/0165-1838(82)90086-8 (1982).

68 Panneton, W. M. et al. Activation of brainstem neurons by underwater diving in the rat. Front Physiol 3, 111, doi:10.3389/fphys.2012.00111 (2012).

69 Gorini, C., Jameson, H. S. & Mendelowitz, D. Serotonergic modulation of the trigeminocardiac reflex neurotransmission to cardiac vagal neurons in the nucleus ambiguus. J Neurophysiol 102, 1443–1450, doi:10.1152/jn.00287.2009 (2009).

70 Gorini, C., Philbin, K., Bateman, R. & Mendelowitz, D. Endogenous inhibition of the trigeminally evoked neurotransmission to cardiac vagal neurons by muscarinic acetylcholine receptors. J Neurophysiol 104, 1841–1848, doi:10.1152/jn.00442.2010 (2010).

71 Layland, J., Li, J. M. & Shah, A. M. Role of cyclic GMP-dependent protein kinase in the contractile response to exogenous nitric oxide in rat cardiac myocytes. J Physiol 540, 457–467, doi:10.1113/jphysiol.2001.014126 (2002).

72 Moen, J. M. et al. Overexpression of a Neuronal Type Adenylyl Cyclase (Type 8) in Sinoatrial Node Markedly Impacts Heart Rate and Rhythm. Front Neurosci 13, 615, doi:10.3389/fnins.2019.00615 (2019).

73 Huq, A. J. et al. A Novel Mechanism for Human Cardiac Ankyrin-B Syndrome due to Reciprocal Chromosomal Translocation. Heart Lung Circ 26, 612–618, doi:10.1016/j.hlc.2016.09.013 (2017).

74 Schultz, T. I., Raucci, F. J., Jr. & Salloum, F. N. Cardiovascular Disease in Duchenne Muscular Dystrophy: Overview and Insight Into Novel Therapeutic Targets. JACC Basic Transl Sci 7, 608–625, doi:10.1016/j.jacbts.2021.11.004 (2022).

75 Roh, H. C. et al. Simultaneous Transcriptional and Epigenomic Profiling from Specific Cell Types within Heterogeneous Tissues In Vivo. Cell Rep 18, 1048–1061, doi:10.1016/j.celrep.2016.12.087 (2017).

76 Habib, N. et al. Div-Seq: Single-nucleus RNA-Seq reveals dynamics of rare adult newborn neurons. Science 353, 925–928, doi:10.1126/science.aad7038 (2016).

77 Todd, W. D. et al. Suprachiasmatic VIP neurons are required for normal circadian rhythmicity and comprised of molecularly distinct subpopulations. Nat Commun 11, 4410, doi:10.1038/s41467-020-17197-2 (2020).

78 Stuart, T. et al. Comprehensive Integration of Single-Cell Data. Cell 177, 1888–1902 e1821, doi:10.1016/j.cell.2019.05.031 (2019).

79 Armstrong, G. et al. Uniform Manifold Approximation and Projection (UMAP) Reveals Composite Patterns and Resolves Visualization Artifacts in Microbiome Data. mSystems 6, e0069121, doi:10.1128/mSystems.00691-21 (2021).

80 Wang, F. et al. RNAscope: a novel in situ RNA analysis platform for formalin-fixed, paraffin-embedded tissues. J Mol Diagn 14, 22–29, doi:10.1016/j.jmoldx.2011.08.002 (2012).

81 Hult, E. M., Bingaman, M. J. & Swoap, S. J. A robust diving response in the laboratory mouse. J Comp Physiol B 189, 685–692, doi:10.1007/s00360-019-01237-5 (2019).

