## Supplemental Figures 1-5 for "Molecular Disambiguation of Heart Rate Control by the Nucleus Ambiguus"

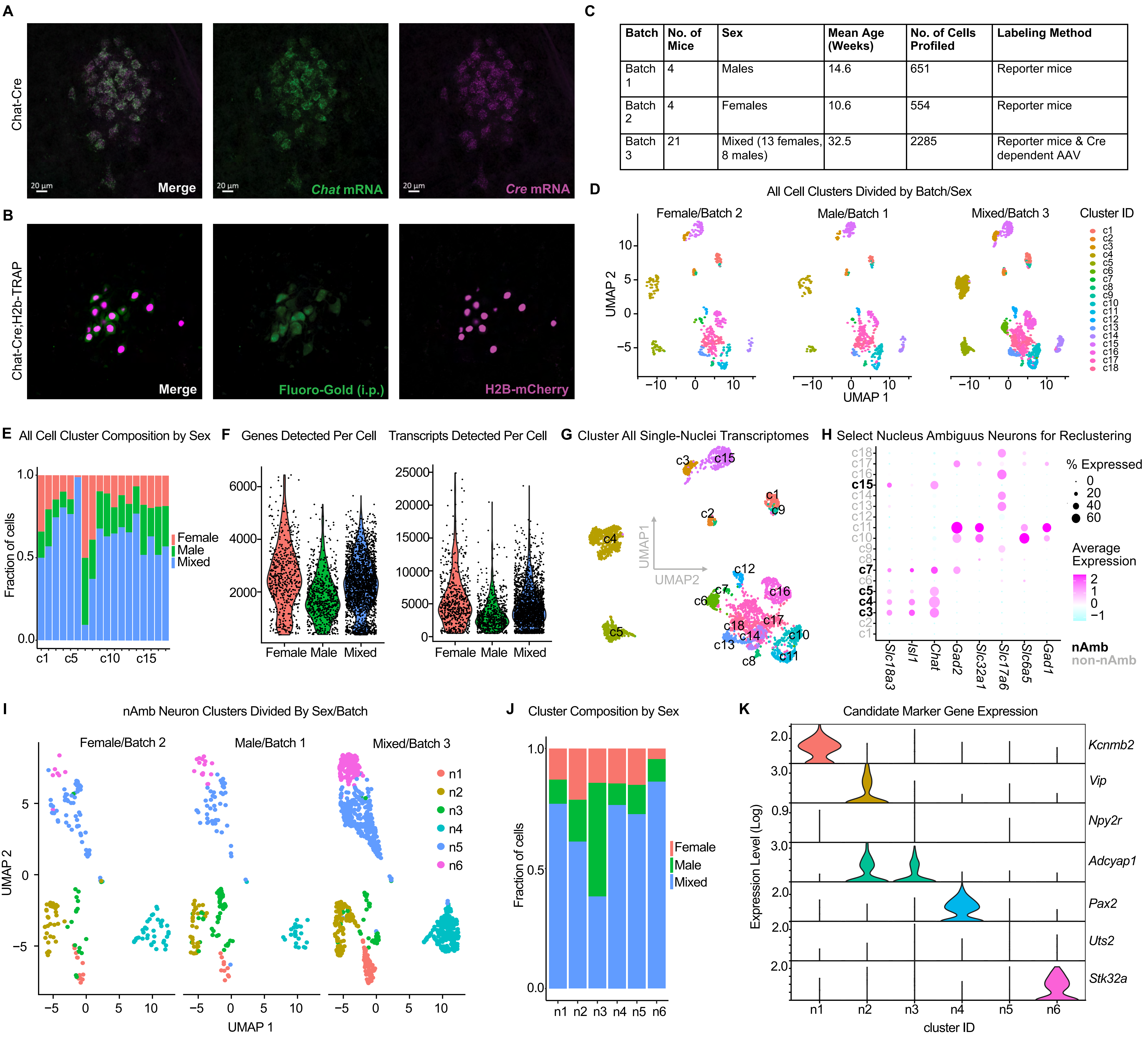


| Batch | No. of Mice | Sex | Mean Age (Weeks) | No. of Cells Profiled | Labeling Method |
|---|---|---|---|---|---|
| Batch 1 | 4 | Males | 14.6 | 651 | Reporter mice |
| Batch 2 | 4 | Females | 10.6 | 554 | Reporter mice |
| Batch 3 | 21 | Mixed (13 females, 8 males) | 32.5 | 2285 | Reporter mice & Cre dependent AAV |

**Supplemental Figure 1: Quality Control of Mouse Lines and snRNA-seq Data**

A. Validation of Chat-Cre expression in the nAmb by RNA fluorescent in situ hybridization (RNA FISH) for *Chat* and *Cre*.
B. Comparison of Chat-Cre;H2b-TRAP labeling in nAmb vagal efferent neurons to labeling of all peripherally-projecting neurons by systemic administration of Fluorogold.
C. Sample metadata for three sample batches of single-nuclei RNA-sequencing data.
D. Batch composition of each cell cluster in all-neuron UMAP.
E. Percent of each all-neuron cluster by batch
F. Numbers of genes and unique transcripts (unique molecular identifiers, UMIs) detected per cell, by batch, in all-neuron dataset.
G. UMAP visualization of all neurons after batch integration.
H. Identification of nAmb neuron clusters based on expression of positive marker genes (*Slc18a3*, *Isl1*, *Chat*) and negative marker genes (*Gad2*, *Slc32a1*, *Slc17a6*, *Slc6a5*, *Gad1*)
I. UMAP visualization of nAmb-only neurons after batch integration.
J. Batch composition of each cell cluster in nAmb-only UMAP.
K. Expression of candidate subtype marker genes in nAmb neuron clusters.

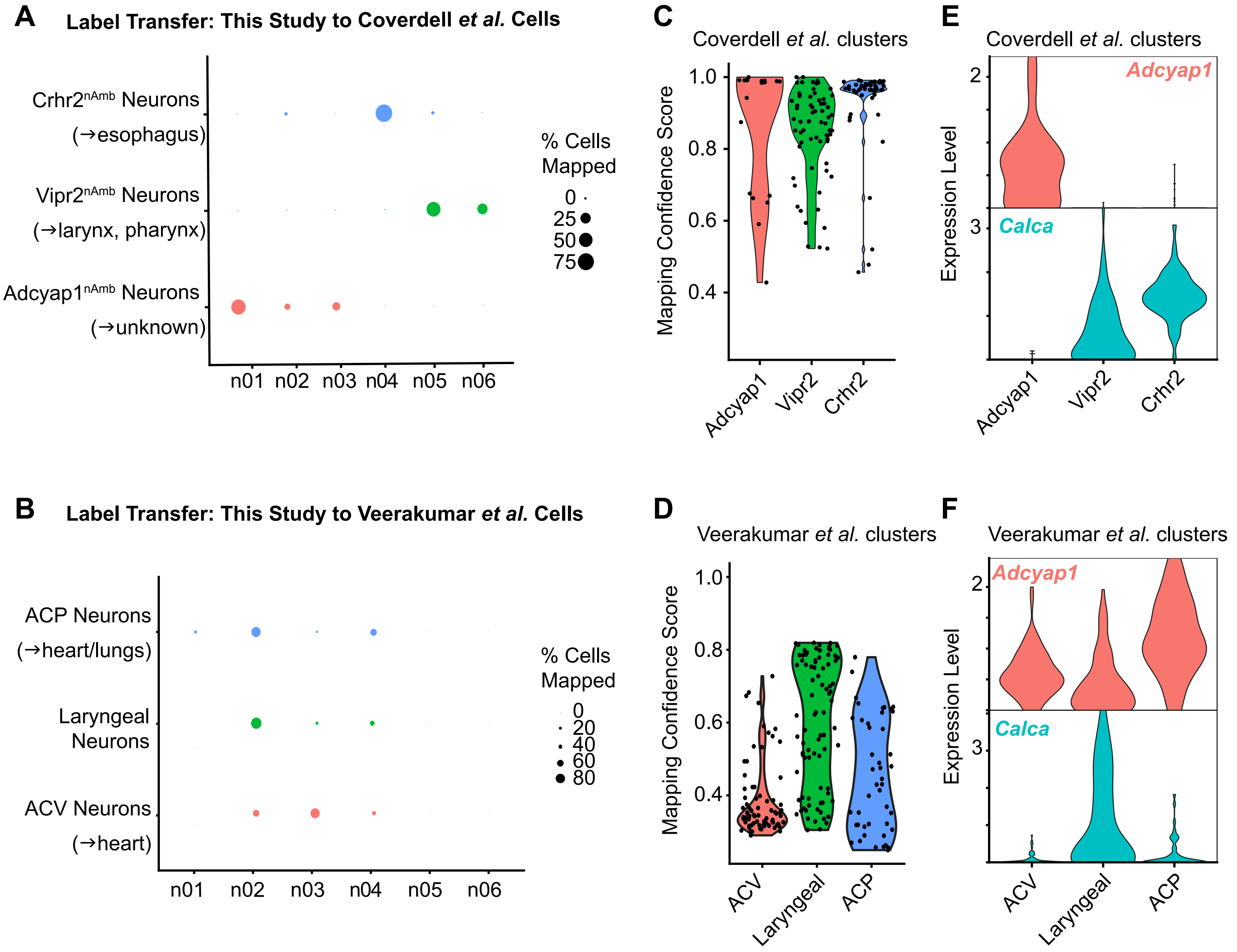

A
Label Transfer: This Study to Coverdell et al. Cells
Crhr2nAmb Neurons
(→esophagus)
Vipr2nAmb Neurons
(→larynx, pharynx)
Adcyap1nAmb Neurons
(→unknown)
n01
n02
n03
n04
n05
n06
% Cells Mapped
0
25
50
75
C
Coverdell et al. clusters
Mapping Confidence Score
1.0
0.8
0.6
0.4
Adcyap1
Vipr2
Crhr2
E
Coverdell et al. clusters
Expression Level
Adcyap1
Calca
2
3
Adcyap1
Vipr2
Crhr2
B
Label Transfer: This Study to Veerakumar et al. Cells
ACP Neurons
(→heart/lungs)
Laryngeal Neurons
ACV Neurons
(→heart)
n01
n02
n03
n04
n05
n06
% Cells Mapped
0
20
40
60
80
D
Veerakumar et al. clusters
Mapping Confidence Score
1.0
0.8
0.6
0.4
ACV
Laryngeal
ACP
F
Veerakumar et al. clusters
Expression Level
Adcyap1
Calca
2
3
ACV
Laryngeal
ACP

**Supplemental Figure 2: Reference Query Mapping**

A,B. Coverdell *et al.* (A) and Veerakumar *et. al* (B) query neuron clusters (y-axis) mapped onto the reference nAmb neuron clusters in the present study (x-axis).

C,D. Confidence of mapping of Coverdell et al. cells (C) and Veerakumar et al. cells (D) onto the nAmb neuron clusters of the current study.

E,F. Expression of *Adcyap1* and *Calca* by nAmb neuron clusters in Coverdell et al. (E) and Veerakumar et al. (F).

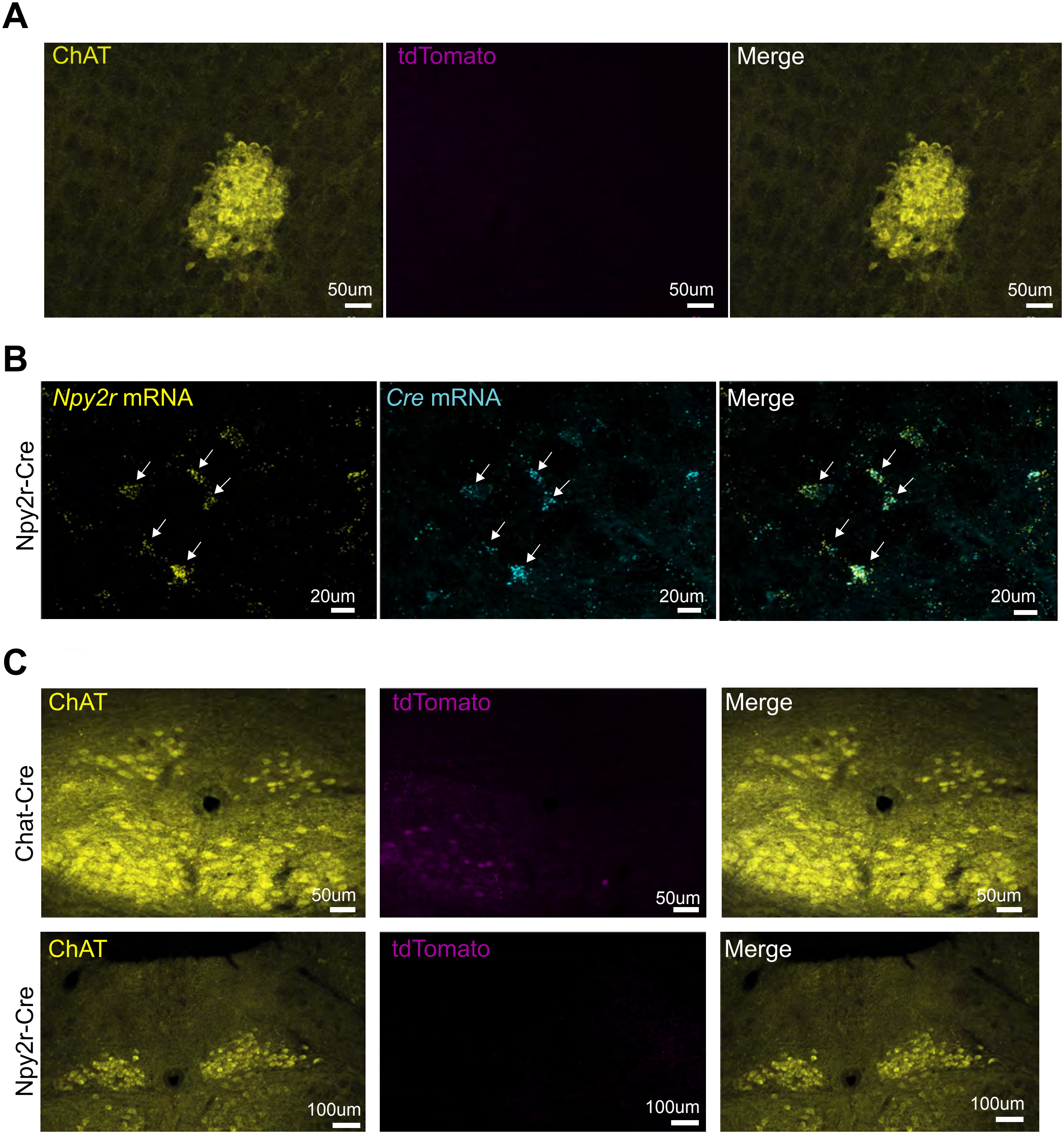
A
ChAT
50um
tdTomato
50um
Merge
50um
B
Npy2r-Cre
Npy2r mRNA
20um
Cre mRNA
20um
Merge
20um
C
Chat-Cre
ChAT
50um
tdTomato
50um
Merge
50um
Npy2r-Cre
ChAT
100um
tdTomato
100um
Merge
100um

**Supplemental Figure 3: Validation of Viruses, Mouse Lines, and Injections**

A. tdTomato expression in C57Bl/6J nAmb after AAV9-DIO-tdTomato injection (negative control). Scale bar, 50um.
B. RNA FISH co-localization *Cre* and *Npy2r* expression in Npy2r-Cre nAmb (n=3 mice).
C. tdTomato expression in dorsal motor nucleus of the vagus (DMV) after AAV-DIO-tdTomato injection into nucleus ambiguus of Cre-expressing mouse lines (n=3 mice per genotype).

Lack of PLAP expression in DMV after nAmb injections

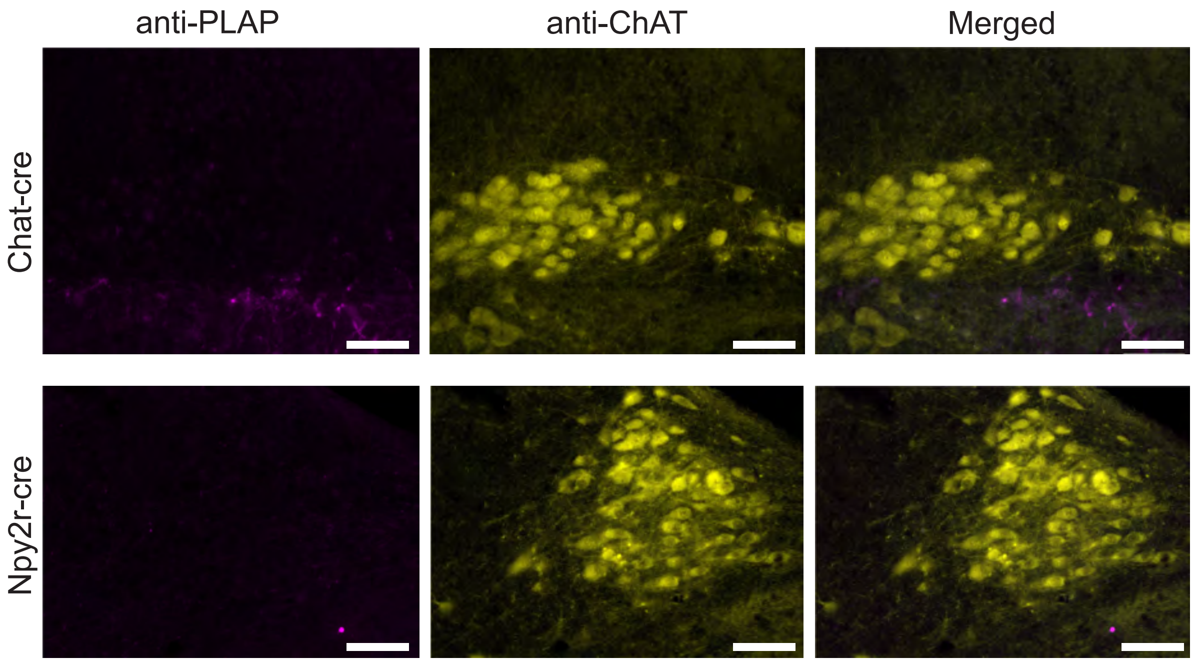

**Supplemental Figure 4: Absence of PLAP-Labeled Cell Bodies in DMV Following nAmb Injections.** No PLAP infected cell bodies were detected in the DMV following injections targeted to the nAmb (n = 3 per genotype). Scale bar: 20 μm.

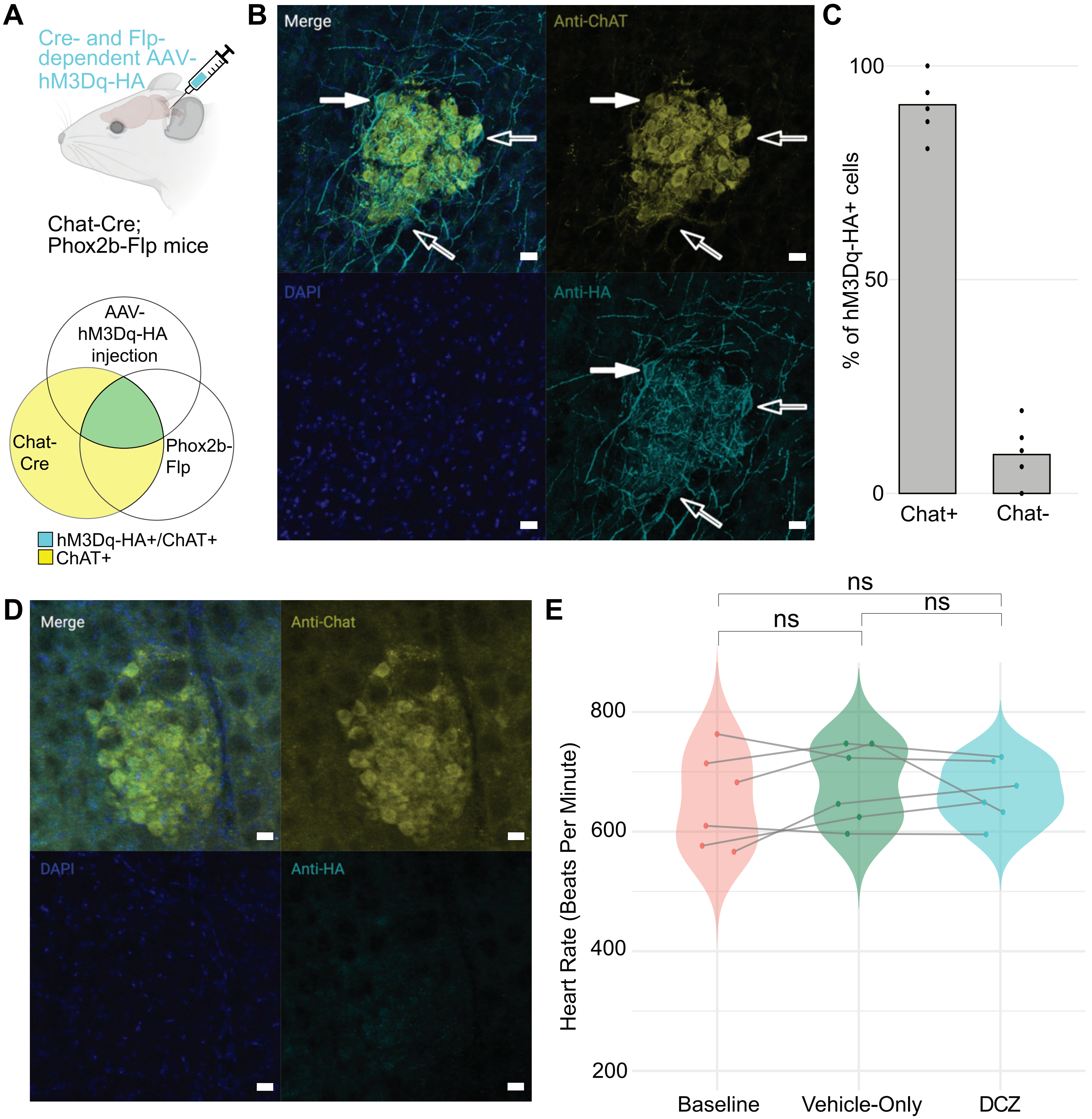
A
Cre- and Flp-dependent AAV-hM3Dq-HA
Chat-Cre; Phox2b-Flp mice
AAV-hM3Dq-HA injection
Chat-Cre
Phox2b-Flp
hM3Dq-HA+/ChAT+
ChAT+
B
Merge
Anti-ChAT
DAPI
Anti-HA
C
% of hM3Dq-HA+ cells
100
50
0
Chat+
Chat-
D
Merge
Anti-Chat
DAPI
Anti-HA
E
ns
ns
ns
Heart Rate (Beats Per Minute)
800
600
400
200
Baseline
Vehicle-Only
DCZ

**Supplemental Figure 5. Validation of Cre- and Flp-Dependent AAV-hM3Dq-HA and Deschloroclozapine.**

A. Schematic of intersectional chemogenetic strategy and expected overlap of hM3Dq-HA and ChAT immunofluorescence.

B. Immunofluorescence of hM3Dq-HA and ChAT after injection of the Cre- and Flp-dependent AAV-hM3Dq-HA into the ventrolateral medulla of Chat-Cre;Phox2b-Flp mice. Solid arrows indicate hM3Dq-HA+ cells with ChAT immunofluorescence, whereas hollow arrows indicate hM3Dq-HA+ cells with little to no ChAT immunofluorescence.

C. Quantification of hM3Dq-HA immunofluorescent cells that were positive or negative for ChAT immunofluorescence

D. Immunofluorescence of hM3Dq-HA and ChAT after injection of the Cre- and Flp-dependent AAV-hM3Dq-HA into the ventrolateral medulla of Phox2b-Flp mice (no Chat-Cre).

E. In Npy2r-Cre animals lacking AAV-hM3Dq, DCZ administration did not alter heart rate ($n = 5$). A one-way repeated-measures ANOVA followed by Tukey's post hoc test showed no significant differences between baseline, vehicle, and DCZ conditions (baseline vs. vehicle: estimate = −28.8 ± 22.6 bpm, $t(10) = -1.28$, adjusted $p = 0.438$; baseline vs. DCZ: estimate = −14.1 ± 22.6 bpm, $t(10) = -0.625$, adjusted $p = 0.810$; vehicle vs. DCZ: estimate = 14.8 ± 22.6 bpm, $t(10) = 0.654$, adjusted $p = 0.794$)
